# Local adaptation to mercury contamination by nitrogen-fixing rhizobia is driven by horizontal gene transfer, copy number, and enhanced gene expression

**DOI:** 10.1101/2023.12.27.573466

**Authors:** Aditi Bhat, Reena Sharma, M. Mercedes Lucas, Kumaran Desigan, Michael Clear, Ankita Mishra, Robert Bowers, Tanja Woyke, Brendan Epstein, Peter Tiffin, José J. Pueyo, Tim Paape

## Abstract

Mercury (Hg) is highly toxic and has the potential to cause severe health problems for foraging animals and humans when transported into edible plant parts. Soil rhizobia that form symbiosis with legumes may possess mechanisms to prevent heavy metal translocation from roots to shoots in plants by exporting metals from nodules or compartmentalizing metal ions inside nodules. Using long-read sequencing, we assembled the genomes of *Sinorhizobium medicae* and *Rhizobium leguminosarum* from the Almadén mercury mine in Spain with high variation in Hg-tolerance to identify structural and transcriptomic differences between low and high Hg-tolerant strains. While independent mercury reductase A (merA) genes are prevalent in α-proteobacteria, Mer operons are rare and often vary in their gene organization. Our analyses identified multiple structurally conserved merA homologs in the genomes of *S. medicae*, including a dihydrolipoamide 2-oxoglutarate dehydrogenase (D2OD), but only the strains that possessed a Mer operon exhibited hypertolerance to Hg. Using RNAseq reads mapped to the unique genome assemblies, we found the Hg-tolerant strains which possessed a Mer operon, that nearly all genes within the operon were significantly up-regulated in response to Hg stress in free-living conditions and in nodules. In both free-living and nodule environments, we found the Hg-tolerant strains with a Mer operon exhibited the fewest number of DEGs in the genome, indicating a rapid and efficient detoxification of Hg^2+^ from the cells that reduced general stress responses to the Hg-treatment. Expression changes in *S. medicae* while inside of nodules showed that both rhizobia strain and host-plant tolerance affected the number of DEGs. Aside from Mer operon genes, *nif* genes which are involved in nitrogenase activity in *S. medicae* showed significant up-regulation in the most Hg-tolerant strain while inside the most Hg-accumulating host-plant, indicating a genotype-by-genotype interaction that influences nitrogen-fixation under stress conditions. Transfer of the Mer operon to non-tolerant strains resulted in an immediate increase in Hg tolerance, indicating that the operon is solely necessary to confer hypertolerance to Hg, despite paralogous merA genes present elsewhere in the genome. This study demonstrated that the Mer operon can be exchanged via horizontal gene transfer into non-tolerant rhizobia strains naturally and experimentally.

## Introduction

Heavy metal stress can have severe consequences for both plant and bacterial cells which exerts strong selective pressure on plant and microbial populations. Mercury (Hg) is a pervasive global pollutant (Driscoll et al., 2013) where the predominant sources of environmental heavy metal contamination, besides natural accumulation or volcanic eruptions, are mines, foundries, and smelters (Tchounwou et al., 2012; Alloway, 2013). Additional sources of heavy metals can be from pesticide and herbicides that that are applied to crops (Rai et al., 2019; Alengebawy et al., 2021). Bioaccumulation in the food chain is commonly in the form of methylmercury (CH_3_Hg^+^), which is highly toxic to humans and other organisms (Poulain and Barkay, 2013; Barkay and Gu, 2021), but inorganic mercury in the form of Hg^2+^ is also highly toxic to most organisms. The main difference between the two lies in their sources as methylmercury, which is most commonly derived from anerobic microorganisms (Parks et al., 2013), and inorganic mercury which typically has an atmospheric or soil deposition source (e.g. mines), or enters through biological systems as a byproduct of demethylation of CH_3_Hg^+^ (Barkay et al., 2003).

Molecular adaptations to mercury stress have been found predominantly in bacteria. The most well characterized genetic mechanism responsible for detoxifying mercury in bacteria is the Mer operon that evolved in geothermal bacteria, but is routinely found in aerobic, anaerobic, aquatic, and terrestrial bacteria as well (Christakis et al., 2021). The Mer operon has the capacity to convert, transport, and detoxify both methylmercury (using MerB) and inorganic mercury (using MerA) along with a suite of membrane transporter genes that are typically tightly linked in the operon (Boyd and Barkay, 2012). Central to the Mer operon is MerA, a flavin-dependent NAD(P)-disulfide (FAD) oxidoreductase that converts Hg^2+^ to Hg^0^, which can be released from the cell as a gas. When the MerA gene is accompanied by MerB, which encodes for an organomercury lyase that catalyzes methylmercury into Hg^2+^ and methane, Hg^2+^ is passed along to MerA where it is transformed to Hg^0^. The MerA and MerB genes were found to be at low frequencies (8% and 2% respectively) among broad taxonomic surveys of bacteria genomes (Christakis et al., 2021), and the difference in frequencies indicate that methyl-mercury detoxification via MerB gene is less prevalent in bacterial genomes than detoxification of ionic mercury. The Mer operon also contains transcriptional regulators such as MerR and MerD, and several proteins that span the inner membrane transport proteins such as MerC, MerE, MerF, MerG, and MerT. In addition to the known Hg ion transporters, glutathione reductase genes (sometimes called MerK, Petrus et al., 2015) can be found in Mer operons as glutathione can potentially transfer Hg^2+^ to MerA (Hong et al., 2010). The overall expression of Mer genes is regulated by MerR, acting as a transcriptional repressor or activator in the absence and presence of Hg^2+^.

Methylation of Hg is primarily the result of microorganisms that use the corrinoid protein, HgcA, and a ferredoxin gene, HgcB (Parks et al., 2013). The benefit of methylation is that it can aid in sequestration of inorganic mercury inside microorganisms, providing a biological mechanism of environmental detoxification of inorganic mercury, but methylmercury is a neurotoxin if it enters animal and human cells. In essence, the HgcAB system in bacteria is responsible for methylation of inorganic mercury, which results in an organo-mercury derivative, CH_3_Hg^+^, while the MerAB system functions to counteract the effects of HgcAB by providing microorganisms the ability to demethylate CH_3_Hg^+^ using MerB, then detoxify inorganic mercury from their cells using MerA. While the Mer operons originated from micro-organisms that survive in geothermal environments (Boyd and Barkay, 2012), MerA and MerB genes are found in broad taxonomic groups. (Christakis et al., 2021) have done the most exhaustive study of the diversity of the MerAB complex in bacteria using phylogenetic approaches of publicly available genomic data and found the presence of these genes in a wide variety of terrestrial bacteria, including soil-borne, nitrogen-fixing rhizobia in the class α-proteobacteria which form symbiosis with plant roots.

Some of the most studied nitrogen-fixing bacteria are *Sinorhizobium medicae* (*syn*. *Ensifer*) and *S. meliloti* due to their symbiotic interactions with the model legume species *Medicago truncatula* and alfalfa (*M. sativa*) (Jones et al., 2007; Reeve et al., 2010a), and *Rhizobium leguminosarum* bv. *trifolii* which mainly interacts with clovers, but also peas (Reeve et al., 2010b; Mendoza-Suárez et al., 2020). While the majority of molecular research in legume host-plants and symbiotic rhizobia has focused on nitrogen fixation mechanisms, which provide obvious benefits to both partners through nitrogen and carbon exchange, more recognition of overlapping stress responses in rhizobia and host plant signaling has been realized to be essential for establishing symbiosis and improved resilience in legume-rhizobia systems (Hawkins and Oresnik, 2022). Symbiotic interactions between legume host plants and nitrogen-fixing rhizobia have profound impacts on soil quality and plant health, and soil microbes in symbiosis with legumes have potential for bioremediation of toxic soils (Fagorzi et al., 2018; Raghupathy and Arunachalam, 2023), particularly if one or both symbiotic partners becomes adapted to a novel stress condition such as heavy metals in the environment.

Tolerance to soils contaminated with toxic heavy metals depends on molecular adaptations that allowed rhizobia to colonize contaminated environments, or adaptations that arose *de novo* through mutation in rhizobia living in the contaminated soils. Because free-living bacteria have such large effective population sizes, adaptive evolution through natural selection on mutations that confer higher tolerance is expected to be strong (Bobay and Ochman, 2017). Moreover, bacterial genome evolution can also occur through horizontal gene transfer (HGT) between strains or between species, providing a source of rapid adaptation to heavy metal stress (Arnold et al., 2022; Chen et al., 2023). Indeed, nitrogen-fixing rhizobia strains isolated from toxic mine sites have revealed they adapted to increases in tolerance to heavy metals (Maynaud et al., 2012, 2014; Nonnoi et al., 2012; Lu et al., 2016). Transcriptomics studies have identified candidate genes that respond to the predominant heavy metals in mining sites by enhanced ion transport mechanisms (e.g., ATPase efflux activity using cadA), but genome wide studies in rhizobia have been mostly focused on adaptations to Cd, Cu, Pb and Zn (Rossbach et al., 2008; Maynaud et al., 2013; Lu et al., 2017; Fagorzi et al., 2018). Studies of adaptive evolution and tolerance to Hg in rhizobia have been limited to a small number of genes (Raghupathy and Arunachalam, 2023), rather than whole genome evolution or genome-wide responses in the transcriptome.

The Almadén mine region in Spain contains some of the greatest concentrations of cinnabar and mercury anywhere on Earth (Garcia Gomez et al., 2006). A large collection of *Sinorhizobium medicae* and *Rhizobium leguminosarum* isolated from root nodules of *Medicago* and *Trifolium* host plants growing at Almadén was measured for tolerance to various heavy metals (Nonnoi et al., 2012). Minimum inhibitory concentrations (MIC) showed that the most Hg-tolerant strains were 10-fold higher than non-Hg-tolerant strains from Almadén, and the increase in tolerance was attributed to *cis*-regulation of merA1 and merA2 (Arregui et al., 2021), the two mercury reductase A homologs not associated with a Mer operon in the genome. The Arregui et al., (2021) study used qPCR to quantify the regulatory changes in two rhizobia strains with different tolerance to Hg, focusing on merA in response to mercury in free-living conditions and in nodules. However, the genomes of these strains were not sequenced and other potential candidate genes or loci that contributed to hypertolerance to Hg could not be identified.

In α-proteobacteria, the class containing the most studied nitrogen-fixing bacteria including the genera *Bradyrhizobium*, *Mesorhizobium*, *Rhizobium*, and *Sinorhizobium*, all possess merA genes. However, most of the sequenced strains lack a Mer operon and these merA genes exist in the genome independent of any Mer operon. In this paper, we will capitalize genes on the Mer operon (e.g., MerA) and homologous mercury reductase A genes will be indicated using lowercase gene names (e.g., merA) to distinguish the difference between the operon and non-operon homologs. The independent merA genes likely contribute to some basal level of Hg tolerance, but non-operon merA genes are unlikely able to confer hypertolerance, which we believe requires a Mer operon in rhizobia. Furthermore, comparisons of protein structures of MerA and merA homologs in rhizobia genomes could provide insight into the conservation of function and whether the homologs provide redundant or elevated tolerance due to additive or multiplied gene expression in genomes that possess MerA, merA1 and merA2.

In addition to heavy metal detoxification responses in rhizobia, it will also be important to identify regulatory changes in nitrogen fixation genes (e.g. *nif*) that would potentially limit nitrogen export to host-plants during symbiosis if these genes are impacted by heavy metals (Jach et al., 2022). Expression of the *nif* gene cluster by rhizobia controls the production of nitrogenase, the necessary enzyme for converting atmospheric nitrogen N_2_ to ammonia (NH_3_) or nitrate (NO_3-_), that can be used as a macronutrient by the host plant. In nodules, the environment for rhizobia changes from purely aerobic in their free-living conditions to a much more anaerobic, oxygen limited environment, inside of nodules. Because heavy metal stress is known to trigger oxidative stress responses in soil microbes including rhizobia (Tan et al., 2010; Jozefczak et al., 2012; Sharma et al., 2012; Maynaud et al., 2013; Alvarez-Rivera et al., 2022), and in *Medicago truncatula* nodules which contain symbiotic *Sinorhizobium* (Villar et al., 2020), heavy metal stress on rhizobia may be even more severe during symbiosis due to the combined metal and oxygen stresses (Bobik et al., 2006).

The Almadén rhizobia strains provide a unique opportunity for comparative genomics and transcriptomics due to known quantitative differences in their tolerance to Hg and other heavy metals. We assembled the genomes of nine *S. medicae* and six *R. leguminosarum* strains from the Almadén mine, using PacBio long-read sequencing to identify structural differences between non-tolerant and tolerant strains. Using the *S. medicae* genomes for paired-end RNAseq read mapping, we used transcriptomics to identify gene expression differences between classes of strains varying in their Hg tolerance in both free-living conditions and inside of nodules. The genome assemblies and differential expression results indicated the presence of complete Mer operons in Hg-tolerant strains, which we claim is the main determinant of Hg hypertolerance, despite the presence of generic merA genes in all *Sinorhizobium* and *Rhizobium* genomes. To test this hypothesis, we transferred a plasmid containing the Mer operon from the most tolerant *S. medicae* strain to two non-tolerant *S. medicae* strains and our findings demonstrated that HGT can instantly confer tolerance to Hg.

## Results

### The mercury tolerant and non-tolerant strains were assembled into complete chromosomes and plasmids

The genomes of the *S. medicae* strains were each assembled into 3-6 contigs with one exception (SupplementaryData1, Table S1), and the *R. leguminosarum* genomes into 6-11 contigs. In most cases these contigs constituted complete circularized chromosomes and plasmids. Three of the Almadén strains (*S. medicae* strains AMp08, SMp01 and *R. leguminosarum* strain STf07) each had a mercury reductase (Mer) operon which was previously unknown in this collection of rhizobia, despite observed high-tolerance to Hg for these three strains in prior studies (Nonnoi et al., 2012). The genome sizes and contig assemblies are consistent with previously published reference genomes of both species, which show *S. medicae* possessing one main chromosome and two symbiotic plasmids (pSymA and pSymB) which contain the symbiotic signaling genes (Geddes et al., 2021), and often an additional accessory plasmid that is unique to individual strains (Reeve et al., 2010a; Ghosh et al., 2021). *R. leguminosarum* tends to have a single large chromosome like *S. medicae,* but typically more plasmids (Young et al., 2006; Reeve et al., 2010a). We performed a phylogenomic analysis using 7 of the Almadén genomes with 65 additional genomes from the Integrated Microbial Genomes and Microbiomes (IMG) database (https://img.jgi.doe.gov) and found that four of the Almadén genomes were located in clades containing *S. medicae* and *S. meliloti,* and the other three strains in an adjacent clade containing *R. leguminosarum* (Figure S1). The closest related strain to the Almadén strains is *Mesorhizobium* sp. LSJC277A00 which contains a Mer operon, but data showing its high Hg-tolerance could not be found. Therefore, the genome assemblies would allow us to make valuable insights into the evolution of local adaptation to Hg tolerance in the Almadén rhizobia strains.

The genomes of the two most Hg-tolerant *S. medicae* strains, AMp08 and SMp01 were assembled into 4 and 6 contigs respectively. We aligned these two focal genomes to the reference *S. medicae* strain WSM419 (Reeve et al., 2010a) using Sibelia to examine synteny and to assign each contig to chromosomes and symbiotic plasmids. In the case of the Hg-tolerant strain AMp08, the main chromosome and two symbiotic plasmids (pSym) are syntenic with the main chromosome and symbiotic plasmids of *S. medicae* WSM419: one chromosome (∼3.8 Mb), pSymA (∼1.2 Mb), pSymB (∼1.6 Mb). It also included an accessory plasmid (∼71 kb) that was not syntenic or shared any homology with the accessory plasmid of WSM419 (Figure 1a). The accessory plasmid in AMp08 which contained a complete mercury reductase (Mer) operon, (Figure S2) is likely derived from a HGT from another rhizobia strain or bacteria species and contains many genes of unknown function (hypothetical proteins).

**Figure 1.**
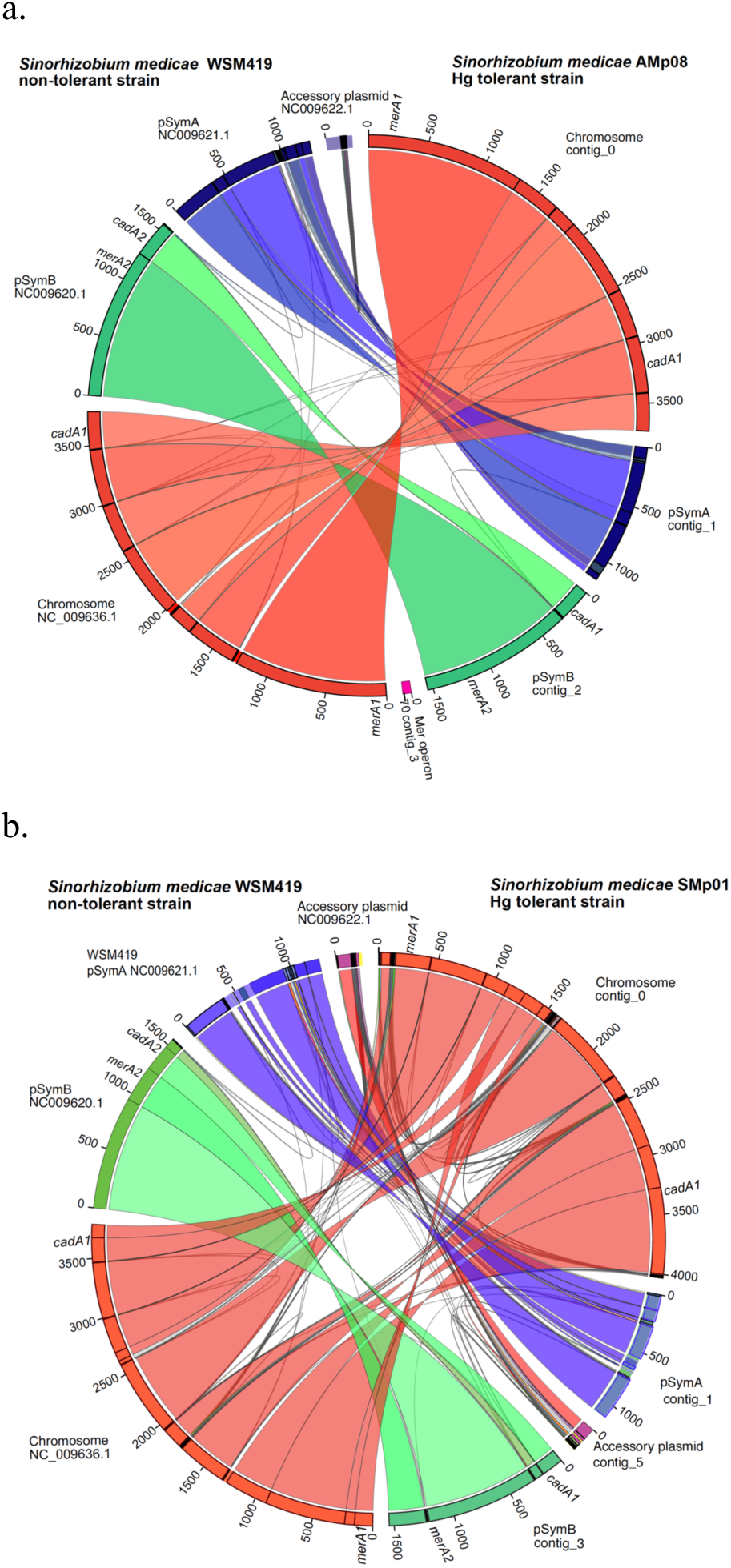
(a) Comparison of the Almadén mine *Sinorhizobium medicae* strain AMp08 (the most Hg tolerant strain which contains a mercury reductase (Mer) operon on an accessory plasmid) with the model *S. medicae* strain WSM419. (b) Comparison of the Almadén mine *S. medicae* strain SMp01 (the most second most Hg-tolerant strain which contains a Mer operon on the main chromosome) with the model *S. medicae* strain WSM419. On the outer ring, common colors were used for homologous chromosomes between the two strains (e.g., red = chromosome, blue = pSymA, green = pSymB), and syntenic regions within chromosomes or plasmids spanning the center of the figure share common colors between the two strains.

We also checked the homology and synteny of the main chromosome and two pSym plasmids of Hg-tolerant *S. medicae* strain SMp01 against WSM419, which showed that 4 of the 6 contigs could be merged (contig_0 with contig_2 and contig_3 with contig_4) resulting in 4 contigs total (Figure 1b). Similar to AMp08, the SMp01 strain also has a Mer operon, but at a different genomic location. In SMp01, the Mer operon is located on the main chromosome and not on the pSym or accessory plasmids, and this region is not syntenic with WSM419 as it lacks the Mer operon. Finally, we aligned the Hg-tolerant *R. leguminosarum* strain STf07 against the *R. leguminosarum* reference strain WSM1325 (Reeve et al., 2010b) and found that this genome was also assembled into complete chromosomes and plasmids that aligned with the reference genome, where both strains possessed one large chromosome and five plasmids (Figure S3). Strain STf07 also possesses a Mer operon, which is located on a smaller plasmid and not on the main chromosome. We therefore have high quality reference genomes for the three most Hg-tolerant rhizobia strains in our collection, each with Mer operons.

### Phylogenetic analysis of mercury reductase A and synteny of the Mer operon in the Almadén strains and diverse species of α-proteobacteria

The three most Hg-tolerant strains among our assembled genomes from Almadén (*S. medicae* AMp08, SMp01 and *R. leguminosarum* STf07) had minimum inhibitory concentrations (MIC) ranging from 200-250µM, while non-tolerant strains typically had MIC of 25 µM (Nonnoi et al., 2012), which is a 10-fold difference in Hg-tolerance. Because mercury reductase A genes (merA1 and merA2) appear to be pervasive in *Sinorhizobium* and *Rhizobium*, we conducted a phylogenetic analysis to identify whether the merA homologs from the highly tolerant strains formed separate clades from the non-tolerant strains, including non-Almadén strains *S. medicae* (WSM419), *S. meliloti* (1021) and *R. leguminosarum* (WSM1325), and other outgroup bacteria species. The phylogenetic analysis of merA homologs in the *S. medicae* and *R. legumonisarum* Almadén strains showed four clades (Figure 2). One clade comprised of mercury reductase A genes that had the greatest sequence diversity, and only included the three tolerant strains (*S. medicae* AMp08, SMp01 and *R. leguminosarum* STf07) along with other outgroup MerA sequences from non-Almadén rhizobia. This clade comprised of MerA genes that were part of a Mer operon, while the merA genes from less tolerant rhizobia strains that lack a Mer operon were not present in this clade and formed separate clades.

**Figure 2.**
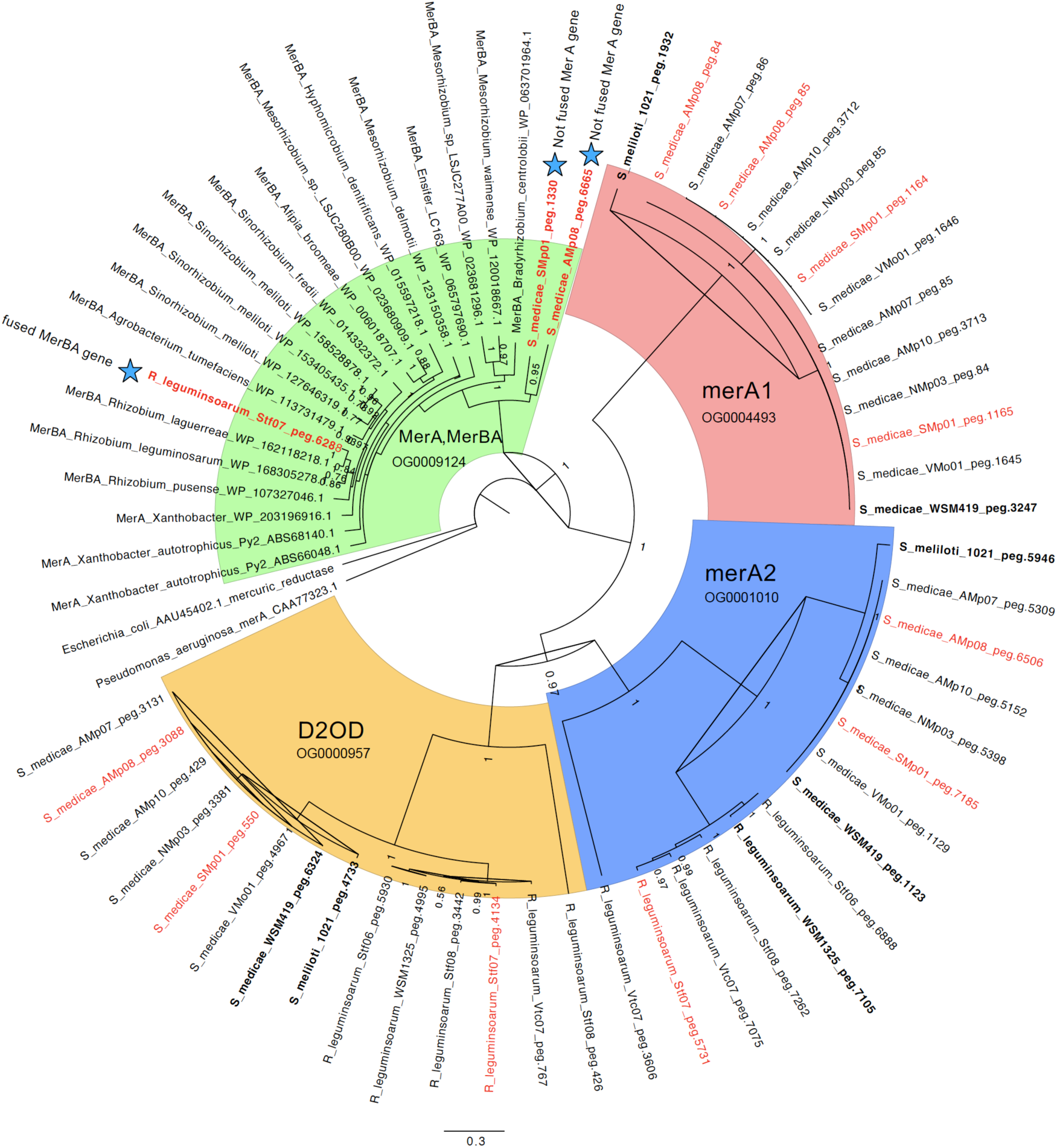
Phylogeny of mercury reductase A (merA) homologs with a focus on *Sinorhizobium medicae* and *Rhizobium leguminosarum*. The merA homologs from the Almadén mine strains that have high tolerance to Hg are indicated by red text. The phylogeny is colored according to four main clades which includes their homolog name and Orthofinder ID. The green clade includes MerA and fused MerBA genes from multiple species/strains of *Mesorhizobium*, *Sinorhizobium* and *Rhizobium* and various bacteria including *Pseudomonas aeruginosa* and *Escherichia coli* (used to root the phylogeny), *Xanthobacter autotrophicus* which has well studied MerA and MerB genes (Petrus et al., 2015), *Agrobacterium tumefaciens, Hyphomicrobion dentrificans, Alfipia broomeae,* and multiple species/strains of *Mesorhizobium*, *Sinorhizobium* and *Rhizobium*. In the green clade are the MerA genes from the Almadén *S. medicae* strains AMp08, SMp01 and *R. leguminosarum* strain STf07 are marked with a blue star. The strain STf07 has a fused MerA and MerB (“MerBA”) (OG0009124) while the strains AMp08 and SMp01 do not have fused MerA and MerB. The merA1 (OG0004493), merA2 (OG0001010), and dihydrolipoamide 2-oxoglutarate dehydrogenase (D2OD) (OG0000957) are colored red, blue and orange, respectively. The numbers on the nodes are posterior probability scores from 500,000 sampled trees using Mr. Bayes v3.2.7 on the CIPRES webserver (https://www.phylo.org/index.php).

Fusions of the MerA and MerB proteins have been found in bacteria where the N-terminal region of MerA was replaced by the alkylmercury lyase domain of MerB (Petrus et al., 2015; Barkay and Gu, 2021). Blast searches of the MerA gene from the Mer operon in the Hg-tolerant *S. medicae* strains against a-proteobacteria and outgroup bacteria including *Pseudomonas*, *Xanthobacter*, and *E. coli* resulted in hits that were often annotated as “MerBA”, despite many lacking an actual fusion of MerA and MerB genes. In many cases, this appears to be an annotation artifact where true fusions of MerA and MerB were classified as MerBA during annotation of deposited genomes due to their close homology to the MerA region of the fused MerBA sequences in GenBank. Among the three most Hg-tolerant strains from Almadén that possessed a Mer operon, one had a MerA gene but no MerB gene (AMp08), one had a MerA and three MerB genes (SMp01), and one had a fused MerBA gene (STf07). Therefore, we conducted a synteny analysis of the Mer operon from the three most Hg-tolerant Almadén strains along with other a-proteobacteria strains available in IMG. The analysis revealed three independent origins of a Mer operon, with each of the Hg-tolerant strains having a Mer operon being more closely related to another species than to eachother (Figure S4), despite all three strains being collected from Almadén. These results indicate that the Mer operons were derived through different HGT sources. Several a-proteobacteria genomes such as *Mesorhizobium* sp. LSJC277A00 had a complete Mer operon, as well as a fused MerBA, but no publications or data that supported a functional Mer operon were found to be associated with these genomes deposited in GenBank or IMG. However, the strains with a complete operon presumably have elevated tolerance to Hg.

In addition to the clade containing MerA homologs that were in Mer operons, our phylogenetic analysis revealed three additional clades. A second clade contained merA1 from all *S. medicae* strains from Almadén, including a short tandem duplication only found in the Almadén strains (merA1’), along with merA1 from *S. medicae* WSM419 and *S. meliloti* 1021 (Figure 2). The merA1 copies from the strains from Almadén and the non-Almadén reference strains showed almost no amino acid sequence diversity among all strains. This suggests merA1 is under strong selective constraint in *Sinorhizobium*, presumably because it provides basic tolerance to Hg^2+^ ions in the environment, regardless of environmental contamination level. A third clade, which we designated as merA2 (annotated as FAD-dependent NAD(P)-disulphide oxidoreductase) based on Arregui et al. (2021), contains all *S. medicae* and *R. legumonisarum* strains derived from Almadén, as well as the non-Almadén strains including *S. medicae* WSM419, *S. meliloti* 1021 and *R. legumonisarum* WSM1325 (Figure 2). *R. legumonisarum* does not possess any homologs in the merA1 clade but has homologs in the merA2 clade. The merA2 gene in this species likely provides the same basic tolerance to Hg^2+^ as does merA1 and merA2 in both *S. medicae* and *S. meliloti*.

A fourth clade which contains dihydrolipoamide 2-oxoglutarate dehydrogenase (abbreviated as D2OD), is a close homolog of merA1 and merA2. An examination of the expression of D2OD in response to Hg-stress may help to determine whether this homologous gene plays a role in Hg detoxification in rhizobia. Finally, because of the independent assemblies and annotations of each of the genomes studied here, we used OrthoFinder to identify homologous genes between each of the genomes. Each of the four genes (MerA/MerBA, merA1, merA2, D2OD) were assigned their own orthogroup which was consistent with their separate phylogenetic clades using Mr. Bayes (Figure 2). We will examine the expression of genes in *S. medicae* represented in the four clades in strains with different tolerance to Hg in later sections of this paper.

### Models of protein structure of mercury reductase A homologs in *S. medicae* show high conservation and the presence of a MerBA fusion protein in *R. leguminosarum*

To estimate whether any structural differences were obvious among the homologous merA proteins (merA1, merA2, D2OD and MerA), we used AlphaFold to create models of each of the homologs and then compared them by superimposing all four structures on each other (Figure 3). To this end, we used amino acid sequences of the four genes from *S. medicae* strain SMp01, which contains a Mer operon. The average pLDDT (ranging from 93 to 96) and PTM score (ranging from 0.92 to 0.94) in AlphaFold (Mirdita et al., 2022) indicated the structural models are highly reliable. The four modelled structures were highly homologous (RMSDs range between 1.0 -1.4 Å) and each one formed a homodimer (Figure 3). The structures revealed the N-terminal FAD-binding domain active site CXXXXC motif, the NADPH-binding domain, the central domain, and the C-terminal dimer interface domain, were all conserved between the four homologs. Previous studies have demonstrated that the N-terminal domain (NmerA) of MerA is responsible for recruiting Hg^2+^ and transferring it to the cysteine pair located at the flexible C-terminal segments. The C-terminal cysteine pair undergoes a conformational change to deliver Hg^2+^ to the active site of the opposing monomer (Ledwidge et al., 2005; Baek et al., 2020). However, in the *S. medicae* MerA homologs, while the N-terminal NmerA domain is absent, the overall structural fold and expected reductase activity appear similar to the catalytic core of the *Bacillus* sp. strain RC607 (Schiering et al., 1991), *P. aeruginosa* (Ledwidge et al., 2005), *L. sphaericus* strain G1 (Bafana et al., 2017), and FDR family proteins (Argyrou and Blanchard, 2004; Baek et al., 2020; Shearer et al., 2023).

**Figure 3.**
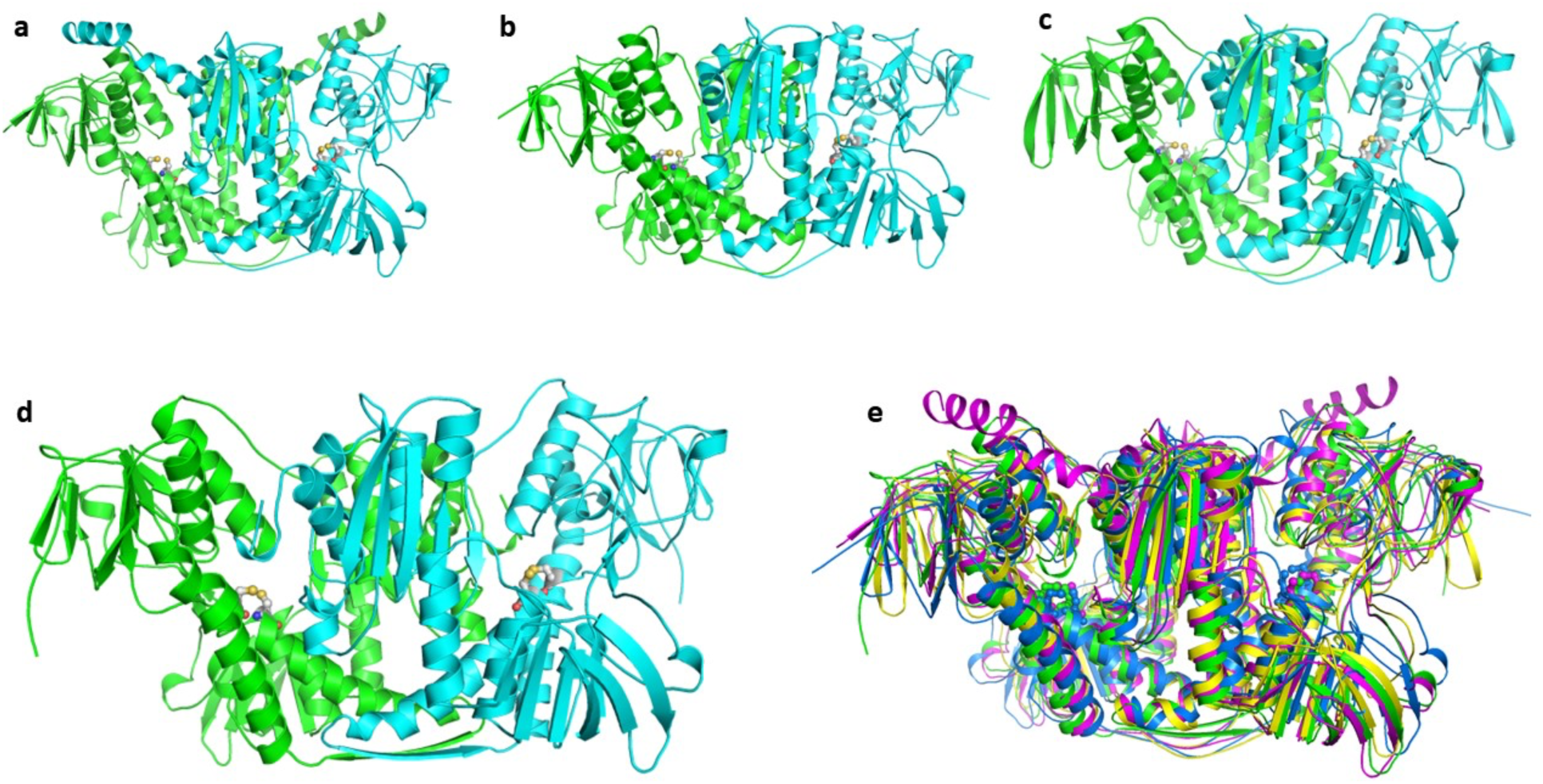
Ribbon representations of the dimeric protein structures of homologous mercuric reductases from *Sinorhizobium medicae* strain SMp01 modeled using AlphaFold. The generic mercury reductase proteins merA1(a), merA2 (b) and D2OD (c) are found in all *S. medicae* and *S. melioti* genomes regardless of their tolerance (Figure 2). The MerA protein (d) from the Mer operon is only found in the Hg-tolerant strains from Almadén (and other publicly available genomes). The two monomers are shown in green and cyan colors. Superposition of structures of merA1 (magenta), merA2 (blue), D2OD (yellow) and MerA (green) (e) show high levels of conservation and homology. Conserved active site cysteine residues are shown as stick models.

To uncover the domain architecture of MerBA fusion proteins in *R. leguminosarum* strain STf07, we modelled the structure of MerBA in this strain and compared it with *Mesorhizobium* sp. LSJC277A00 using AlphaFold (Figure S5). The structures of MerBA from both *R. leguminosarum* strain STf07 and *Mesorhizobium* sp. LSJC277A00 were homodimeric and the MerA component was highly homologous to the non-fused MerA proteins from *S. medicae.* In both protein structures, the MerB domain is tethered to MerA by a flexible linker and minimally interacts with N-terminal FAD-binding domain of MerA (Figure S5c). The structure of the MerB domain was found to be similar to *E. coli* (Lafrance-Vanasse et al., 2009) and the conserved active site residues Cys 147, Cys 212 and Asp 150 were located at the central core of the domain. To our knowledge, this is the first reported protein structure for a fused MerBA protein, at least in nitrogen-fixing a-proteobacteria.

### Differentially expressed genes (DEGs) in response to Hg stress in free-living conditions are mostly Mer operon genes in highly tolerant rhizobia

We selected four *S. medicae* strains with variable tolerance to Hg and treated them with 4 μM HgCl_2_ in free-living conditions for transcriptomic analysis (Supplementary Data 2). We found that the most tolerant strains had dramatically fewer differentially expressed genes (DEGs) when compared to non-tolerant strains (Figure 4a). The two tolerant strains AMp08 and SMp01, had only 17 (0.2% of the total genome) and 32 (0.5% of the total genome) DEGs respectively, which were mostly upregulated (60%-90%). The low Hg-tolerant strains AMp07 and VMo01 had 540 (7.9% of the total genome) and 448 DEGs (10.3% of the total genome) respectively. This amounts to roughly 25 times more DEGs in the sensitive strains than in tolerant strains, and there were almost equal proportions of upregulated and downregulated genes (40-55%) in the non-tolerant strains. These strain-specific patterns can be easily visualized by comparing the volcano plots (Figure 3 b-d, OrthoFinder IDs were used to label genes due to the independent assembly of each reference genome), which also show key genes involved in Hg detoxification such as MerA (OG0007479), MerT (OG0006346), MerB (OG0010667), and MerP (OG0005882) among the other most significant DEGs (Supplementary Data 2). The very low number of DEGs, and mostly upregulated genes in Hg-tolerant strains, suggests quick triggering of the Mer operon gene expression and highly efficient removal of Hg from the cells. On the other hand, the higher number of DEGs in low tolerant strains, and more equal ratios of up to down-regulated genes, suggests greater cellular and genomic impact of Hg stress on the susceptible strains.

**Figure 4.**
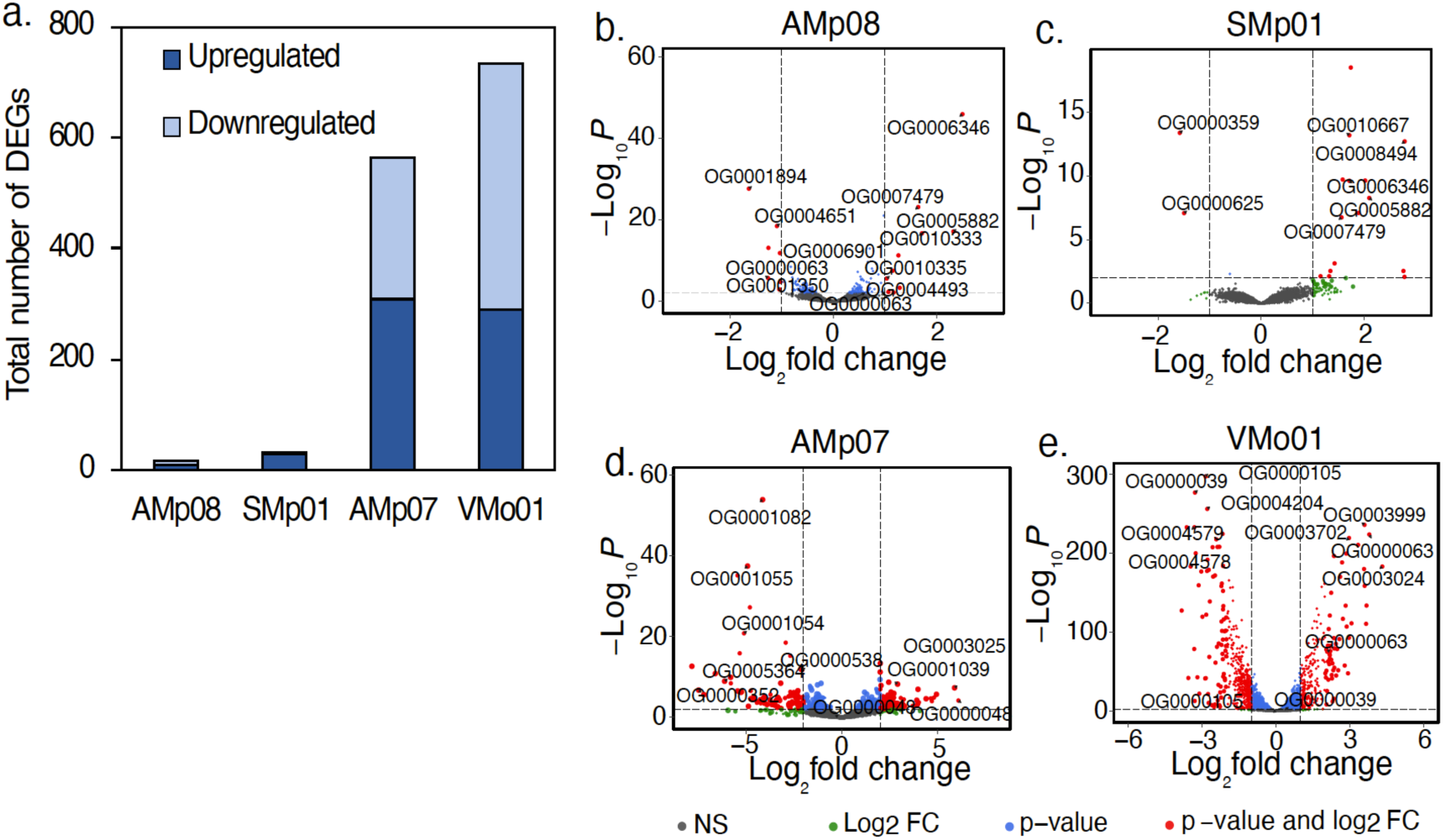
Differentially expressed genes (DEGs) in free-living conditions following Hg stress (a). Free-living rhizobia were grown in the absence (0 μM, Control) or presence of 4 μM HgCl_2_ (Hg-treated). DEGs were identified based on the criteria |log_2_FC| ≥ 1, adjusted p < 0.05, and those that met these criteria are shown in red in the volcano plots for AMp08 (b), SMp01 (c), AMp07 (d) and VMo01 (e). The strains AMp08 (b) and SMp01 (c) each have a complete mercury reductase operon (see Figure 2). Each point represents a gene, and the top significantly upregulated and downregulated DEGs were labeled using their respective Orthogroup IDs.

For the non-tolerant strains AMp07 and VMo01 which lack a Mer operon, several genes from the cytochrome O-ubiquinol oxidase gene family (OG0001054, OG0001055) and nitric oxide dioxygenase (OG0001082) were significantly downregulated, and genes from ABC-transporter families such as ABC-transporter substrate-binding protein (OG0003025), ferric-ion iron-binding ABC-transporter protein (OG0003702), and beta-galactoside, ABC-transporter substrate-binding protein (OG0003024, OG0003025) were significantly upregulated (Figure 4d, 4e; Supplementary Data 1, Table S2). These genes and gene families are known to be involved in mediating oxidative stress responses and ABC-transport mechanisms of cellular compartmentalization of toxic ions (Prabhakaran et al., 2016), which can be expected to be more responsive in the absence of an effective mercury detoxification mechanism such as a Mer operon.

### Significant up-regulation of mercury reductase operon (Mer) genes in Hg-tolerant strains but not in the generic merA genes

As reported above, many of the significant DEGs that responded to Hg stress belonged to the Mer operon. While the generic mercury reductase genes merA1 and merA2 are present in all the *S. medicae* strains as stand-alone genes, only the Hg-tolerant strains possess the Mer operon (*S. medicae* strains AMp08 and SMp01) which includes a MerA gene surrounded by other genes that transport and detoxify Hg in the operon locus. In the two most Hg-tolerant strains, AMp08 and SMp01, a majority of the DEGs, including MerA, showed significant up-regulation in response to Hg. In the Mer operon, we also observed significant upregulation for MerT (mercuric transport protein), MerB (organomercurial lyase, all three copies in Smp01), MerC (cellular membrane transport), MerP (periplasmic mercury transport), MerF and other non-characterized hypothetical proteins (Figures 5 a, b). It is noteworthy that upregulation of MerB occurred despite no methyl-mercury (CH_3_Hg^+^) present in our culture media, indicating that ionic Hg is enough to trigger upregulation of all genes in the Mer operon. Interestingly, we did not find any significant expression change of the transcriptional regulator MerR in the Mer operon in either strain, suggesting constitutive expression of MerR.

**Figure 5.**
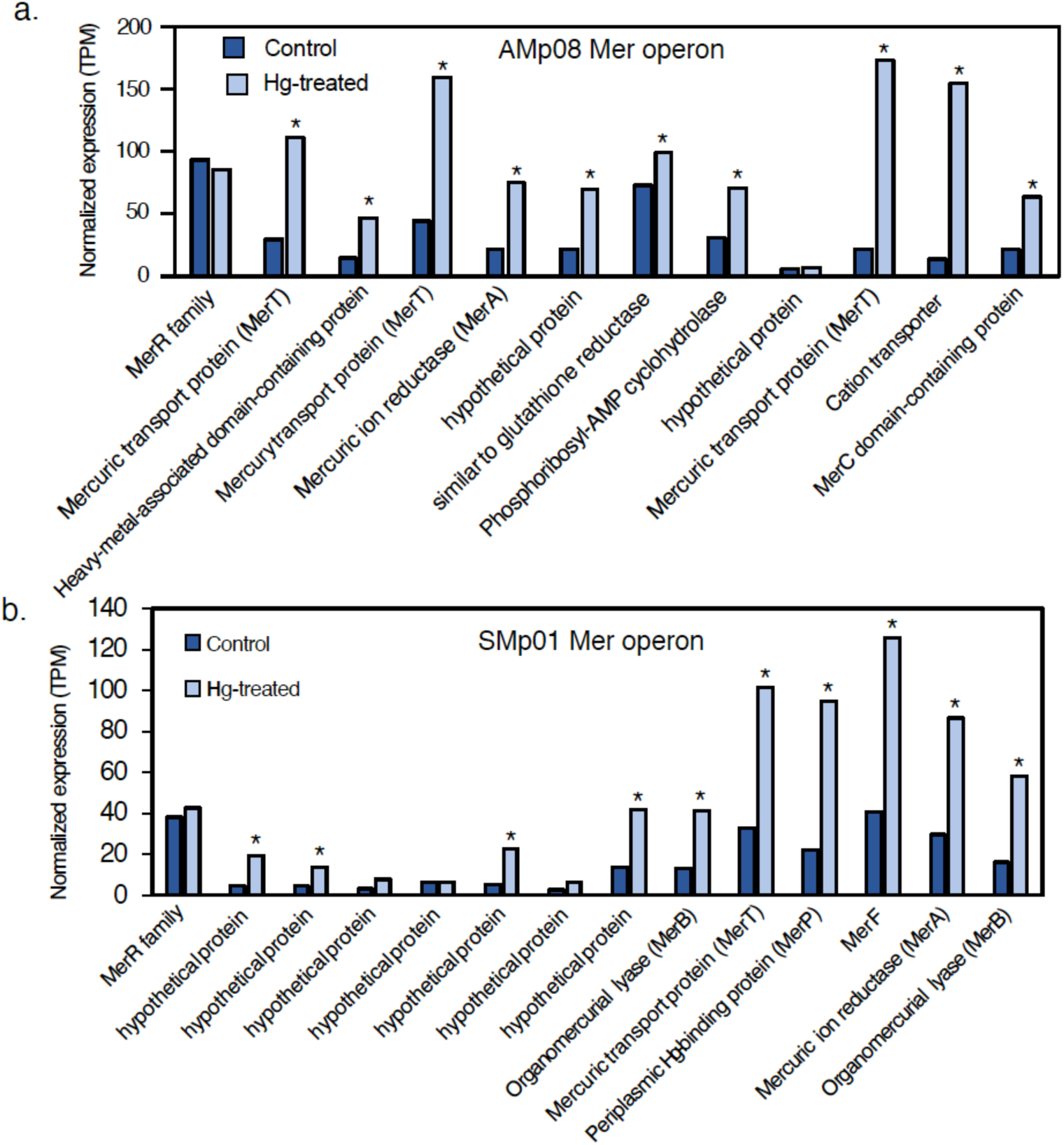
Expression of Mer operon genes in AMp08 (a) and SMp01 (b) in control conditions compared with Hg treated. The strain AMp08 does not possess a MerB gene. In SMp01 MerB is present in two copies. Free-living rhizobia were grown without Hg (Control) or presence of 4 μM HgCl_2_ (Hg-treated). Three biological replicate cell cultures were grown for control and Hg-treated. Normalized expression levels are reported as the mean TPM of three biological replicates.

The common mercury reductase gene merA1 (OG0004493), which transforms Hg^2+^ into the volatile form Hg^0^, is present in all the strains analyzed in the current study regardless of Hg-tolerance. We observed upregulation of merA1 in response to Hg stress in the Almadén strains, however the change was not significant in neither tolerant nor susceptible strains (Figure 6a). Similarly, another mercury reductase annotated as FAD-dependent NAD(P)-disulphide oxidoreductase, referred to as merA2 (OG0001010) (Arregui et al., 2021), showed non-significant changes in expression (Figure 6b). D2OD (OG0000957), which is homologous to the merA genes (Figure 2), showed a significant upregulation in response to Hg treatment in the non-tolerant strains only (Figure 6c). It is unclear whether this gene contributes to some level of Hg tolerance but an extensive search of merA homologs by Boyd and Barkay (2012) across a broad range of bacteria also found dihydrolipoamide dehydrogenases to be homologous to merA, but experimental evidence of their role in Hg tolerance or detoxification is lacking.

**Figure 6.**
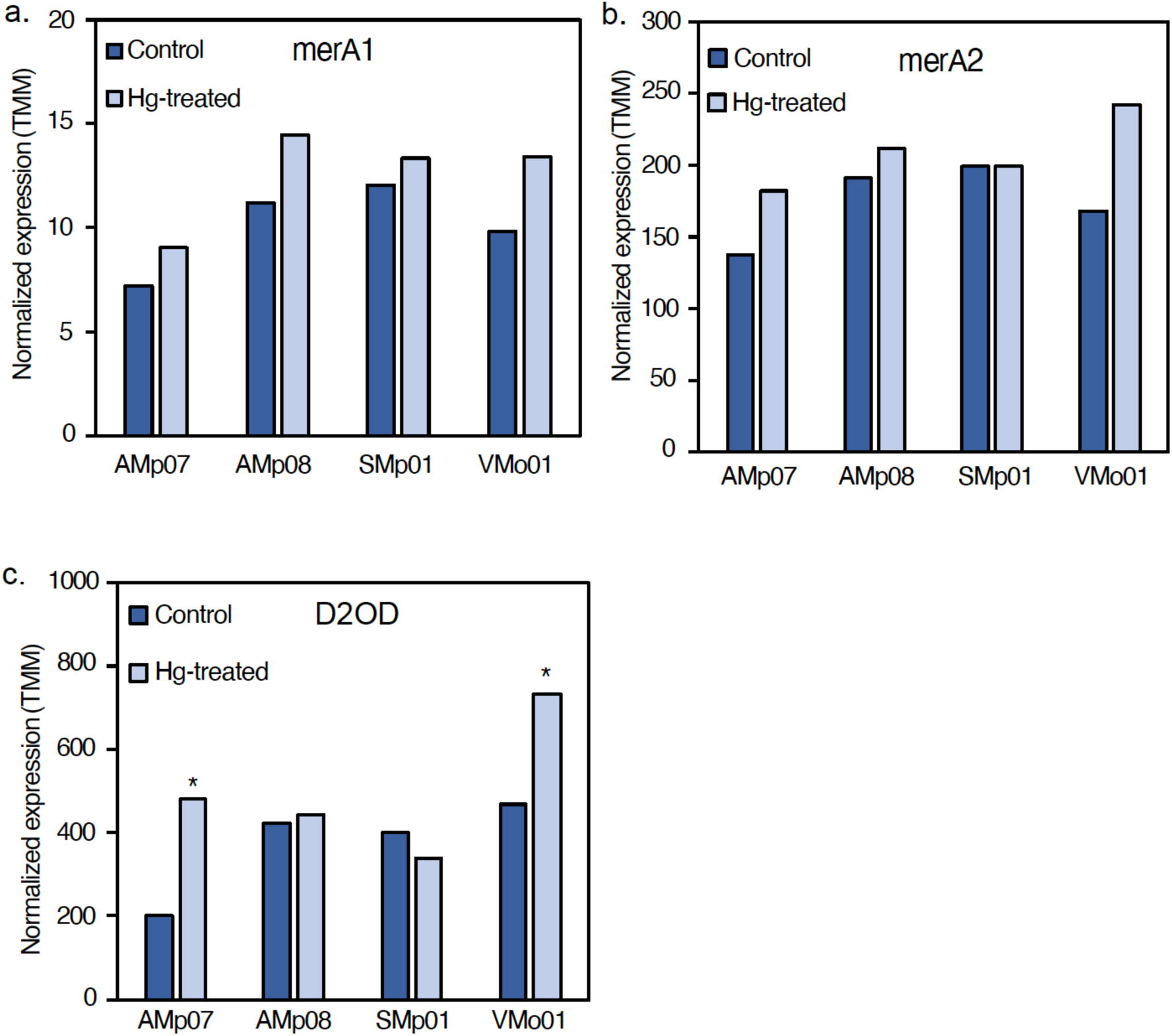
Changes in expression of mercuric ion reductase gene homologs in different strains following Hg treatment. (a) Mercury ion reductase (merA1) showed no significant change in response to Hg treatment in any of the strains studied, although the highest expression was found in AMp08. (b) Mercury ion reductase (merA2) shows no significant change in response to Hg stress in any of the strains tested. All three of these genes, merA1, merA2 and D2OD are homologous to the MerA gene in Figures 2 and Figure 5 (c) A gene annotated as a Dihydrolipoamide 2-oxoglutarate dehydrogenase (D2OD), which is homologous to merA1 and merA2 and shows a significant upregulation in non-tolerant strains (AMp07 and VMo01), and no significant changes in the tolerant strains (AMp08, SMp01). The normalized expression levels are reported as mean TMM values of three biological replicates.

### Comparison of DEGs in response to Hg stress in nodules shows the effect of symbiosis and genotype x genotype interactions

To test the effect of Hg stress on rhizobia in their symbiotic state inside of host-plant nodules, we performed an experiment using two *M. truncatula* host-plant genotypes (high Hg-accumulating genotype HM304 and a low Hg-accumulating genotype HM302 (Paape et al., 2022)) inoculated with either a Hg tolerant (AMp08) or non-tolerant (AMp07) *S. medicae* strain. Following 21 days post-inoculation, the plants were treated with 100 μM HgCl_2_. The purpose was two-fold: the first to provide a biologically relevant rhizobia response as the nodule is a drastically different environment than free-living conditions, and second, there is a possibility of genotype-by-genotype interactions between the rhizobia strains and the host plants that may be observed due to both partners having variation in tolerance or accumulation of Hg. As the RNA isolated from the nodule is a mixture of both the host-plant and rhizobia, we focused on the rhizobia responses in the current study, and the plant responses will be reported separately (Sharma et al. *in preparation*).

We found a stark difference between the number of DEGs in the nodules formed by the tolerant strain AMp08 in the nodules of both host-plant genotypes when compared to the non-tolerant strain, AMp07. In the nodules, AMp08 showed 186 genes and 330 genes in the host-plants HM304 and HM302 respectively (Figure 7a). By comparison, AMp07 had 873 DEGs in HM304 while those in HM302 had 639 DEGs, which was a 1.8 and 4.5-fold increase in the number of DEGs in AMp07 compared with AMp08 in the two host-plants (Figure 7a), respectively. These results show a similar trend with what we observed under free-living conditions where the tolerant strains also showed lower numbers of DEGs than the non-tolerant strains. However, compared with the free-living rhizobia conditions, there was an overall higher proportion of DEGs in the genome when in nodules, as 3-12% of the genome was differentially expressed when in nodules, compared to only 0.2-10% of the genome under free-living conditions. The higher proportion of rhizobia genes responding in nodules may be due to a stronger stress treatment in the experiment conducted *in planta,* or the presence of other environmental factors inside the nodules, including signaling between host-plants and rhizobia.

**Figure 7.**
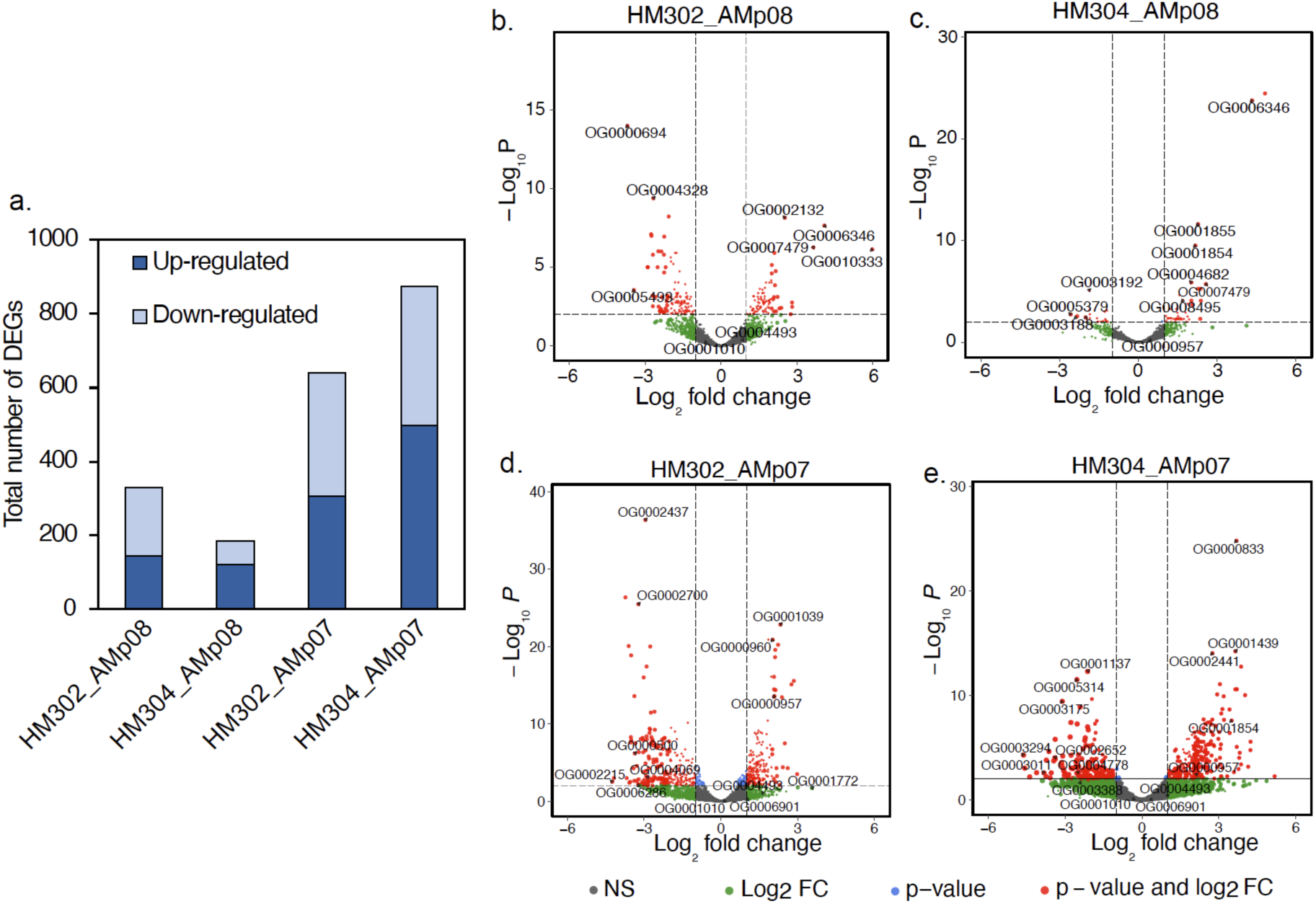
Total number of differentially expressed genes (DEGs) for rhizobia present in nodules following Hg treatment (a). The host-plants inoculated with rhizobia were treated with either 0 μM HgCl_2_ (Control) or 100 μM HgCl_2_ (Hg-treated). DEGs were calculated based on the criteria |log_2_FC| ≥ 1, adjusted p < 0.05. Volcano plots for DEGs for host plants inoculated with AMp08 (HM302_AMp08 (b), HM304_AMp08(c)), and host plants inoculated with AMp07 (HM302_AMp07 (d), and HM304_AMp07 (e)). Each point on the volcano plot represents a DEG, and the top significantly upregulated and downregulated DEGs were labeled using their respective Orthogroup IDs.

For the Hg-tolerant strain AMp08 inside nodules, similar to the free-living conditions, we found the majority of DEGs were located on the Mer operon including MerA, MerT, and glutathione reductase, which had a 3-12-fold increase in expression levels in response to Hg stress in both host-plant genotypes (Figure S6). The total number of shared DEGs between free-living conditions and inside of nodules for the tolerant strain AMp08 amounted to only 0.6% of the DEGs (3 genes total), which included MerA and MerT, indicating the critical role of Mer operon genes in Hg detoxification in free-living conditions and nodules. The merA1 and merA2, which are not present on the operon, showed different patterns than in free-living conditions where merA1 had greater upregulation but was not significant (Figure S7). For the non-tolerant strain AMp07, 2.8% of the DEGs were common between free-living conditions and the nodules. The shared DEGs included multiple genes from the ABC-transporter family which are known to play a role in metal ion transport and cellular detoxification in plants and bacteria. We found that the identity of most of the DEGs under free-living conditions and in nodules is largely different, which could be due to many factors including the cellular replication of rhizobia inside the nodules, signaling between host-plant and rhizobia, and the dilution of the stress by some contribution from the host-plant response. But overall, the response of the genes in the *S. medicae* Mer operon appear to be playing the key role in Hg detoxification based on their up-regulation patterns in the nodules, which is a similar trend in free-living conditions (Figure 5, Figure S6).

When globally examining the most significant DEGs in strain AMp08 (with lowest adjusted p-values shown in Figure 7), we found certain genes such as polyribonucleotide nucleotidyltransferase to be highly upregulated in rhizobia when inside the HM302 host-plant, while phosphoserine aminotransferase and D-3-phosphoglycerate dehydrogenase were significantly upregulated in the HM304 host-plant (Figure 7 b, c; Supplementary Data 1, Table S3). For the AMp07 strain inside HM302 nodules, we found significant downregulation of NAD-dependent protein deacetylase (SIR2 family), manganese catalase, and ABC-transporter substrate-binding protein, while genes such as Na^+^/H^+^-dicarboxylate symporter, malate dehydrogenase and dihydrolipoamide dehydrogenase of 2-oxoglutarate dehydrogenase were significantly upregulated (Figure 7d). Inside of nodules of the HM304 host-plant, AMp07 genes that were differentially expressed included ATP synthase F0 sector subunit a, nitric oxide reductase activation protein, D-3-phosphoglycerate dehydrogenase, all of which were upregulated, while glutathione S-transferase, kup system potassium uptake protein, and ABC-transporter permease protein were downregulated (Figure 7e). These results indicate there are expression patterns that are unique to the rhizobia strains in different host-plant environments and responses depend on the level of tolerance to Hg by both the rhizobia strain and the host-plant genotype.

To visualize strain specific patterns of differential expression between free-living and nodule environments, we selected the top 10 significantly upregulated and top 10 significantly downregulated genes in each strain, then compared the expression of these 20 genes per strain across all the strains for a total of 160 genes (Figure 8, Supplementary Data 1, Table S4). This allowed us to visually compare clusters of the most differentially expressed genes in each strain for sets of homologous genes in the other strains. For each strain we found distinct clusters of significantly high-and low-expressed genes which were often not differentially expressed in other strains, and typically coincided with their tolerance levels (measured using MIC) of the strain. For the tolerant strains, there was a clear cluster of genes corresponding to the Mer operon being significantly upregulated in both free-living conditions and in nodules, while no such cluster was observed for the downregulated genes.

**Figure 8.**
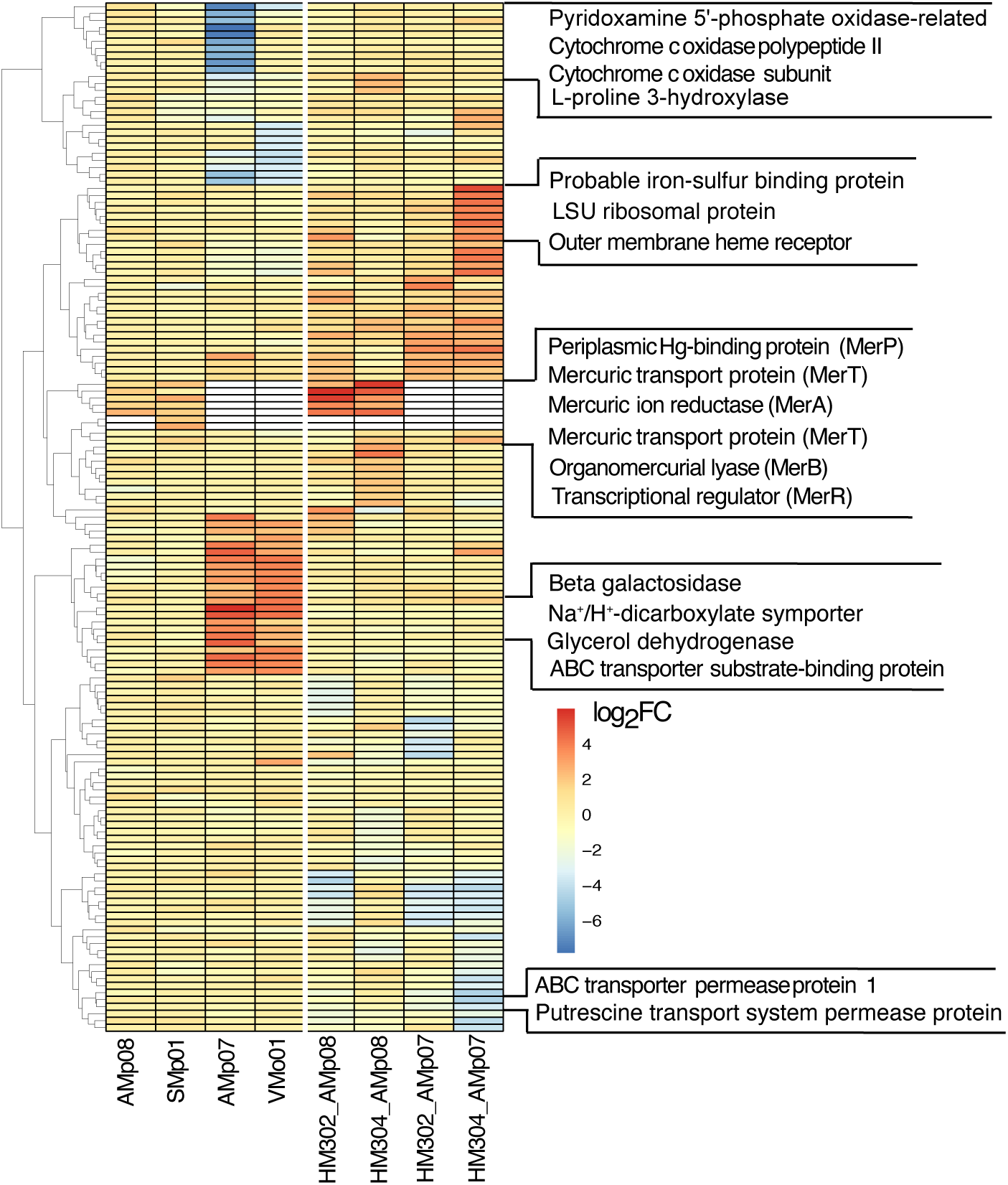
Heatmap representing the top 20 DEGs (top 10 upregulated and top 10 downregulated DEGs) of each sample set for a total of 160 DEGs across all samples encompassing free-living and nodules conditions. On the y-axis, genes and gene families with biological relevance to Hg transport and detoxification were labeled. For the rhizobia strains with no mercuric ion reductase (Mer) operon genes (MerP, MerT, MerA, MerB, MerR), the cells that are white indicate no data point as these strains do not possess a Mer operon or one or more of the Mer operon genes.

For the non-tolerant strains growing in free-living conditions, we observed a cluster of downregulated genes belonging to the family of cytochrome c oxidase subunits as well as genes such as pyridoxamine 5’-phosphate oxidase-related and L-proline 3-hydroxylase. By contrast, clusters of up-regulated genes included transporters such as ABC-transporter substrate-binding proteins, Na^+^/H^+^ dicarboxylate symporter genes, beta galactosidase and glycerol dehydrogenases. Inside nodules, the non-tolerant strain AMp07 had several highly expressed genes including an iron-sulphur binding protein and an outer membrane heme receptor clustering together, while several permeases such as ABC-transporter permease protein and putrescine transport system permease protein formed a distinct cluster of low-expressed genes. Taken together, this analysis allows us to distinguish the responses of Hg-tolerant strains from non-tolerant strains based on the distinct cluster of genes that are differentially expressed in each strain in response to Hg stress, which showed a much broader impact on the transcriptome while in symbiosis than in free-living conditions.

### Hg-stress has an impact on expression of nitrogen fixation (*nif*) genes in symbiosis with host-plants

We conducted GO-enrichment of DEGs from the rhizobia transcriptome in free living conditions and in the nodules to better understand the broader response to Hg stress in *S. medicae* outside of the Mer operon or merA1 and merA2 genes. Overall, GO-enrichment was not very powerful to identify GO-terms in the Hg-tolerant strains as there were too few DEGs to detect enrichment (Supplementary Data 1, Table S5). We found enrichment of the GO-term for nitrogen fixation (GO:0009399) in nodules, which came from differentially expressed nitrogen fixation (*nif*) genes in the HM304-AMp08 host-plant-rhizobia combination. We therefore compared expression of *nif* genes in all four strains grown under free-living conditions, and in the two strains used to inoculate the host-plant to determine whether *nif* regulation was predicted by the tolerance of the strains in free-living or nodule conditions. In free-living conditions, the least tolerant strain AMp07 showed a significant down-regulation of a majority of the *nif* genes (*nifA*, *nifB*, *nifT* and FNA) suggesting a relationship between tolerance and nitrogen-fixation gene expression (Figure S8). In nodules, the strain AMp08 (Hg-tolerant) with the higher Hg-accumulating phenotype host-plant (HM304), showed an upregulation in the *nif* genes in response to Hg treatment, whereas the expression levels of the *nif* genes in AMp08 were downregulated inside the nodules of the low accumulating (HM302) plant genotype (Figure 9 a, c). By contrast for AMp07 (Hg-non-tolerant), we observed a significant downregulation of *nif* genes (*nifA*, *nifB*, *nifT* and FNA) in AMp07 inside the nodules of both host plant genotype (Figure 9 b,d), suggesting that *nif* genes are negatively affected by Hg stress in the non-tolerant rhizobia strain regardless of the host-plant tolerance. The finding that AMp08 *nif* genes were up-regulated inside the HM304 (Hg-high-accumulator) host plant, but down-regulated in the non-tolerant HM302 host plant, suggests a genotype-by-genotype interaction which may allow for better nitrogen fixation by the AMp08 strain in the presence of Hg, if the host plant also possesses higher Hg-stress detoxification capacity. By contrast, for the non-tolerant strain AMp07, the pattern of down-regulation of these genes in both the host-plant genotypes (Figure 9 b, d), and in free-living conditions were highly similar (Figure S8).

**Figure 9.**
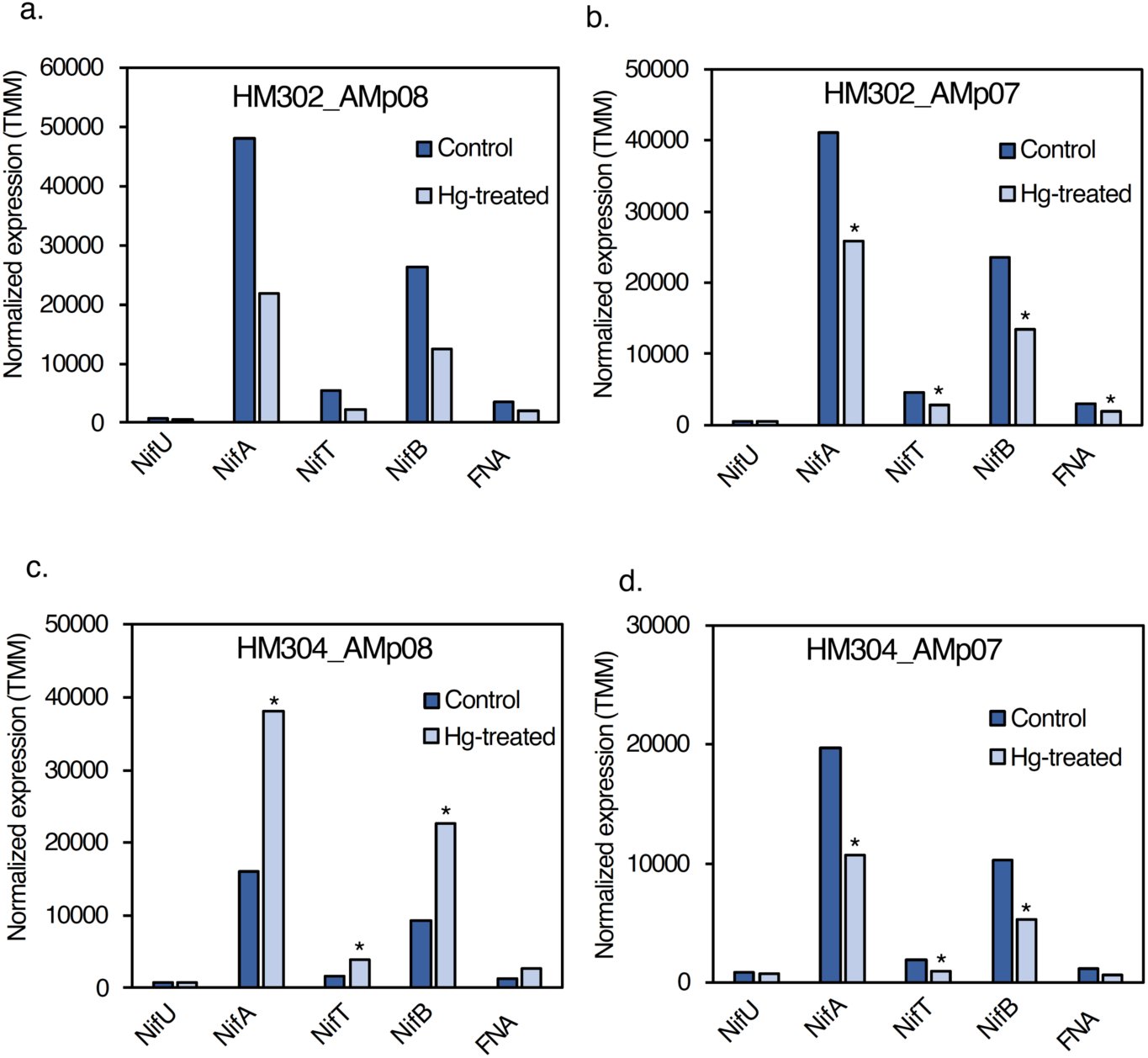
Expression of *nif* genes in rhizobia present in nodules under Hg treatment. In the nodules formed by tolerant strain AMp08, the expression levels of *nifU*, *nifA*, *nifT* and 4Fe-4S ferredoxin nitrogenase-associated gene (FNA) were significantly downregulated when the host plant was HM302 (a), and significantly upregulated when the host-plant was HM304 (b). In the nodules formed by non-tolerant strain AMp07, the expression levels of *nifU*, *nifA, nifT* and 4Fe-4S ferredoxin nitrogenase-associated gene (FNA) were significantly downregulated when the host plant was either HM302 (c) or HM304 (d). The normalized expression levels are shown as the mean TMM values of three biological replicates.

### The Mer operon from Almadén strain AMp08 can confer Hg-tolerance to non-tolerant strains through horizontal gene transfer

Our transcriptomics analysis showed the genes present on the Mer operon were the most significant DEGs in the tolerant AMp08 strain under free-living conditions and in nodules (eg. MerT, MerA). Because all strains contain merA1 and merA2 and show some response to Hg treatments, but probably not enough to confer hypertolerance like in AMp08 or SMp01, we set out to demonstrate that the Mer operon was essential for conferring Hg tolerance. We isolated the 71 kb accessory plasmid from AMp08 (Figure S2) harboring the Mer operon, then transferred the entire plasmid to the two non-tolerant strains AMp07 and WSM419 using electroporation. Without the Mer operon, both the wild-type strains, AMp07 and WSM419 could grow at 25 μM HgCl_2_ (Figure S9), but only the wild-type AMp08 strain grew at greater concentrations. We therefore tested the ability of the transformed strains to grow on higher concentrations of HgCl_2_ than the wild-type strains without the plasmid (Figure 10a). After the Mer operon was transferred into AMp07 and WSM419 strains, they gained the ability to grow at 250 μM HgCl_2_, which is equivalent to the tolerant AMp08 strain (Figure 10a). We confirmed the positive transfer of the Mer operon by amplifying MerA and MerT that are only present on the operon (Figure 10 b-c), while all strains contained merA1 (located on main chromosome) as expected (Figure 10d). To compare the relative expression levels of the Mer genes in the non-transformed and transformed strains grown in the presence or absence of 4 μM HgCl_2_, we performed qPCR (primers listed in Supplementary Data 1, Table S6). While we did not observe any significant change in the expression levels of merA1 and merA2 in any of the strains tested, we found upregulation in the genes present on the Mer operon (MerA, MerT) which had been transferred into AMp07 and WSM419 (Figure 10 e-g), similar to the native Mer operon of the wild-type AMp08 strain. These results confirm that the Mer operon is key to Hg tolerance, and the tolerance can be transferred to other non-tolerant strains via horizontal gene transfer.

**Figure 10.**
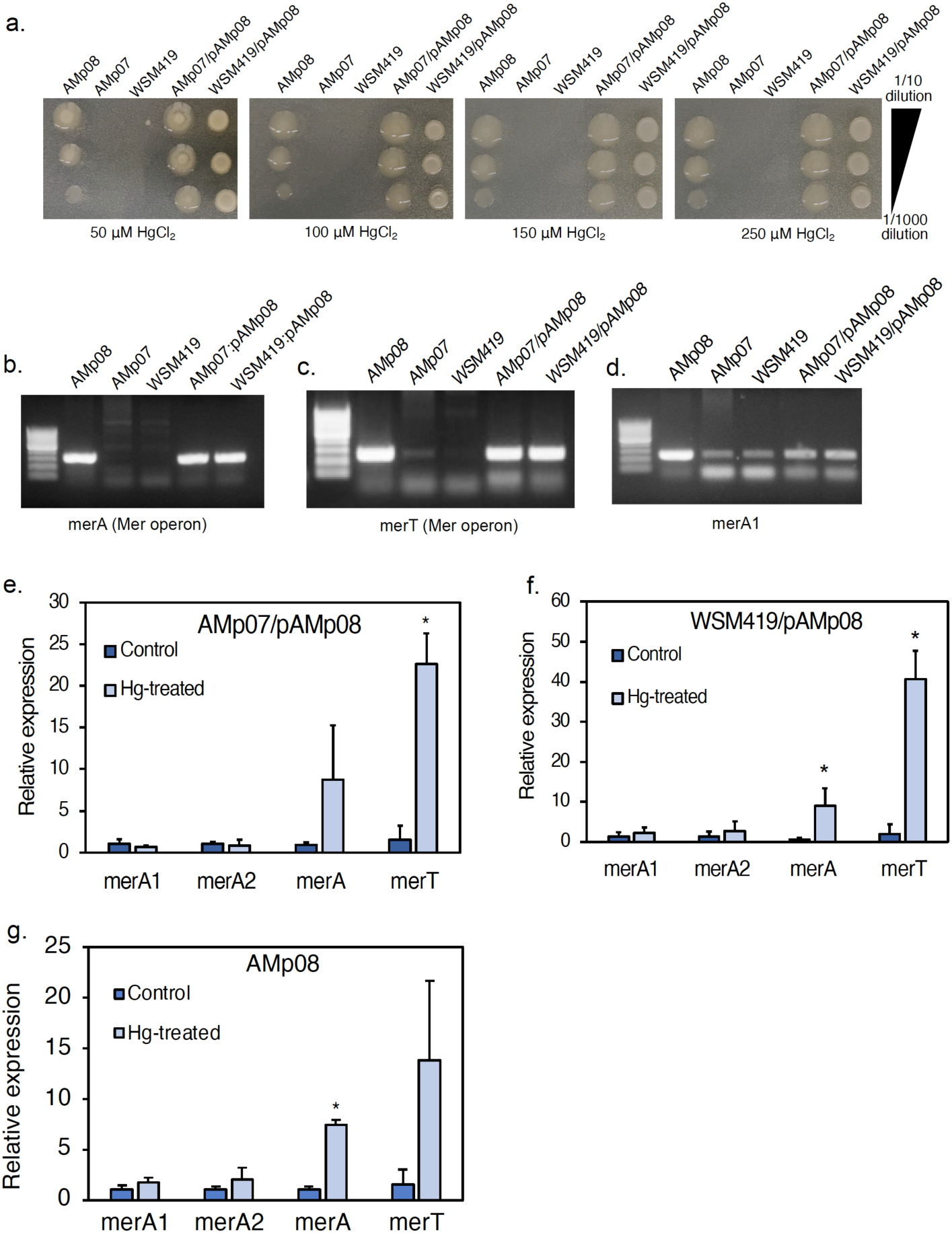
Minimum inhibition concentration (MIC) assay shows the gain of Hg tolerance in non-tolerant strains transformed with plasmid from AMp08 (AMp07/pAMp08; WSM419/pAMp08). Each MIC image represents *S. medicae* strains from left to right: AMp08, AMp07, WSM419, AMp07/pAMp08, WSM419/pAMp08, grown on plates containing 50 μM HgCl_2_, 100 μM HgCl_2_, 150 μM HgCl_2_, 200 μM HgCl_2_, 250 μM HgCl_2_ respectively (a). Each strain was grown at three ten-fold dilutions (from top to bottom): 1/10, 1/100 and 1/1000. Using genomic DNA, PCR was used to amplify the Mer operon genes MerA and MerT, and the generic merA1 gene located on the main chromosome but not on the accessory plasmid containing the Mer operon. Gel images show the presence of MerT and MerA in AMp08 and the transformed strains AMp07/pAMp08, WSM419/pAMp08, but not in the untransformed strains (AMp07, WSM419) that lack a Mer operon (b, c). Gel image showing the presence of merA1 (present on the main chromosome) in all the strains (d). The ladder size scale on each gel image is 50 bp. Expression changes in merA1, merA2, MerA, and MerT in AMp07/pAMp08 and WSM419/pAMp08 quantified using qPCR (e, f). Relative expression levels of merA1 (present on the main chromosome), merA2 (present on pSymB) and MerA, MerT (present on the Mer operon accessory plasmid) were calculated using 16S rRNA as an endogenous control. Three independent experiments were conducted and mean values ± SD are represented. Asterisks indicate p-value < 0.05 calculated using a Student’s T-test.

## Discussion

The genomes we assembled have shown the presence of a complete Mer operon in three of the most Hg-tolerant strains known in rhizobia. The Mer operon constitutes a structural adaptation in these genomes which allowed tolerate high levels of Hg exposure at Almadén. In the *S. medicae* genomes, we identified four mercury reductase A homologs (merA1, merA2, D2OD, MerA) which were all present in the Hg-tolerant strains, while the non-tolerant strains lacked the MerA gene. This new finding advanced our understanding of Hg-tolerance in these strains which was previously attributed to differential expression and regulatory changes in the mercury reductase genes, merA1 and merA2 (Arregui et al., 2021), which are not part of a Mer operon. The synteny and phylogenetic analyses showed that the Mer operons in the three strains were most likely acquired independently, which is a remarkable example of convergent evolution at a local geographic scale. The diverse synteny is also consistent with previous observations of high levels of diversity in gene organization and structural diversity of Mer operons in bacteria in general (Naguib et al., 2018). Recently, it has been shown that varying degrees of metal stress in the local population may increase the frequency of HGT events (Chen et al., 2023), providing a source of adaptive evolution in bacteria (Arnold et al., 2022) in contaminated environments. Moreover, HGT is also crucial in rhizobia to determine their host range (Msaddak et al., 2023). And while we typically expect selection to act on standing genetic variation in a species (Barrett and Schluter, 2008), the mechanism of HGT can cross species boundaries, accelerating adaptation, even if the donor genetic locus is at low frequency in the population (Woods et al., 2020) or in the microbiome of the rhizosphere. To our knowledge, the presence of a functional Mer operon and its relationship to other mercury reductases in a-proteobacteria has yet to be reported, despite the considerable number of genomic studies of nitrogen-fixing bacteria in this group (Sugawara et al., 2013; Nelson et al., 2018), including strains collected from heavy metal contaminated sites (Maynaud et al., 2013, 2014; Lu et al., 2016, 2017).

### Regulatory changes in Hg-tolerant rhizobia in free-living conditions low compared to non-tolerant strains

Transcriptomic analysis showed several interesting patterns that were dependent on the presence of the Mer operon as well as the environmental conditions associated with growth of the rhizobia (i.e., whether free-living or in symbiosis). The number of DEGs in the two Hg-tolerant strains in free-living conditions was remarkably low (∼20-30 genes), while the non-tolerant strains had nearly 25 times more DEGs. Among the most significant DEGs, nearly all genes located in the Mer operon in both Hg-tolerant strains were significantly up-regulated. However, there was an exception, the regulatory gene MerR showed constitutive expression in both control and Hg-treated conditions. The very low number of DEGs suggests the rapid and efficient mechanism of Hg^2+^ transport, volatilization, and elimination from the cell results in extremely low impact on free-living rhizobia. The low number of DEGs in the Hg-tolerant strains appears to be in line with heavy metal tolerant *Mesorhizobium metallidurans* (Maynaud et al., 2013) and *Sinorhizobium meliloti* (Lu et al., 2017), which also showed few genes differentially expressed (∼1.0 % or less of the genome showing significant responses), but these studies did not compare tolerant strains with less tolerant strains from the same population or elsewhere. We suspect that adaptation to high levels of Hg (and other toxic heavy metals) in the soil and rhizosphere is most important in free-living conditions due to direct exposure to the metal ions in the environment. However, we must concede that the level of Hg stress in our free-living experiment (4 μM HgCl_2_) was a low dose that was necessary for obtaining transcriptomic data from both the non-tolerant strains and tolerant strains, but higher doses would potentially result in greater numbers of stress response DEGs in the tolerant strains.

We were surprised to find that the non-operon mercury reductases (merA1 and merA2) showed only slight up-regulation between control vs. treatment conditions and appeared more like constitutively expressed genes and showed only slight differences between tolerant and non-tolerant strains. However, the highest expression of merA1 in Hg-treated conditions was in our most tolerant strain, AMp08, which may provide additional tolerance above what the MerA gene provides. Overall, our findings suggests that merA1 and merA2 in *S. medicae* (and likely *S. meliloti*) function in basic Hg^2+^ detoxification but they cannot be optimized by *cis*-regulation to achieve hypertolerance to the scale that the Mer operon provides, as indicated by the low Hg-tolerant strains from the same population. Moreover, we found almost no amino acid sequence diversity for the merA1 gene and this gene appears fixed in Sinorhizobium, which would fit the HGT followed by gene specific sweep model shown by Arnold et al., (2022), which could mean merA1 was also the product of an earlier HGT event and other Mer genes were lost in environments where high Hg tolerance was unnecessary. Furthermore, the very low diversity for the merA2 gene among the several strains that were included in the phylogenetic analysis (including the two non-Almadén strains *S. medicae* WSM419 and *S. meliloti* 1021), is consistent with strong selective constraint on both merA1 and merA2 in *Sinorhizobium* to confer basic tolerance to Hg (i.e., < 25μM) in free-living conditions. Finally, we speculate that the significant up-regulation of D2OD in the two low Hg-tolerant strains but not in the two most Hg-tolerant strains that possess a Mer operon, may be the result of a local adaptation (i.e., enhanced *cis*-regulation) that may contribute to an inducible mercury reduction process due to high exposure to Hg^2+^ in their native environment at Almadén, but further study into the function of this gene in Hg detoxification is necessary.

### Expression of genes inside of nodules also shows Mer genes dominate the response to Hg stress

Because rhizobia also exist in symbiotic life-stages, it was important to examine gene expression patterns while inside of nodules of legume host plants. It is interesting to find a similar trend of fewer DEGs in the Hg-tolerant strain inside of the nodules, but the total number of DEGs for both tolerant and non-tolerant strains is higher than in free-living conditions. This could be attributed to the higher concentration of the stress treatment (100 μM Hg compared to 4 μM Hg under free-living conditions), but the differences in free-living and nodule life stages cannot be underestimated due to the tremendous signaling between host-plant and rhizobia that occurs to maintain homeostasis during stress responses in nodules. Like in free-living conditions, some of the most significant genes in nodules were those found on the Mer operon. Globally, genome-wide stress responses based on the most differentially expressed genes in the non-tolerant strains revealed many expected types of genes including oxidative stress related genes, cytochrome c oxidases, cytochrome O-ubiquinol oxidase, nitric oxide dioxygenase, ABC-transporters, and other membrane related transporters. The relationship between reactive oxygen species and symbiosis appears to be linked (Hawkins and Oresnik, 2022) and disentangling these responses from the heavy metal stress is challenging.

Due to higher number of DEGs compared with free-living conditions, GO-enrichment of the top ranked DEGs revealed some genes that may be involved in the oxidative stress and cellular detoxification response that resulted from the Hg exposure to the rhizobia in the nodules of the host-plant. The GO-term for nitrogen fixation (GO:0009399) was enriched in nodules, which included several nitrogen fixation (*nif*) genes which regulate nitrogenase levels. The effect of regulatory changes on *nif* genes could provide a more direct link to stress impacts on nitrogen fixation than the more general responses of oxidative stress. It is noteworthy that the least tolerant *S. medicae* strain in our study (AMp07) showed significant down-regulation of *nif* genes in free-living conditions, but the role of the *nif* genes in nitrogenase production and nitrogen-fixation is more relevant inside the host-plant. In a previous study using the AMp08 genotype in *M. truncatula* nodules, Arregui et al., (2021) did not detect any reduction in nitrogenase activity in plants treated with 500 μM Hg when compared with non-treated plants, but the expression levels of neither *nif* nor other nitrogen fixation genes were measured. The *nif* genes in the AMp08 strain showed no significant changes in free-living conditions but in nodules, both the tolerant strain AMp08 and non-tolerant strain AMp07 showed down-regulation while residing in the less Hg-accumulating host plant genotype (HM302). In the higher Hg-accumulating genotype (HM304), AMp08 showed up-regulation but AMp07 showed down-regulation of *nif* genes. Therefore, if up-regulation of *nif* genes is positively correlated with nitrogenase production, then the host-plant genotype x rhizobia strain interaction may also be important for nitrogen fixation under stress conditions. It is noteworthy that the number of nodules is greater in both host-plants when in symbiosis with AMp08 compared with AMp07, but the nodule number was even greater in the HM304 host plant with AMp08 following the stress treatment (Sharma et al, *in preparation*). Because higher nitrogen content in the host plants will improve plant fitness under stress conditions, by proxy the nodule number phenotype could be the result of the rhizobia strain tolerance and its ability to fix nitrogen under Hg stress conditions (Jach et al., 2022).

### Horizontal gene transfer of the Mer operon accessory plasmid achieved hypertolerance in *S. medicae*

Regardless of the role of other genes involved in dealing with Hg stress, our horizontal transfer experiment clearly indicated that an accessory plasmid containing the Mer operon locus alone would suffice to deliver hypertolerance to Hg. This is consistent with functional studies of the Mer operon in other bacteria (Petrus et al., 2015). The gene expression patterns in free-living conditions and inside of nodules of highly up-regulated Mer operon genes, and the immediate acquisition of hypertolerance to Hg when the Mer operon was transferred into non-tolerant strains, provides evidence that the Mer operon is the sole source of this high level of tolerance. MerT, the essential transport protein which facilitates the transport of the Hg^2+^ ion from MerP to MerA (Morby et al., 1995) was one of the highest expressed genes present on the Mer operon in any context, including free-living conditions, nodules and following HGT. MerT is likely one of the most important transporters in the operon, next to MerA and MerB. The Mer operon is unique among heavy metal transport mechanisms in that it can provide a near complete source of cellular detoxification and elimination of the target metal, unlike Cd, Pb or Zn which showed that knocking out several genes is needed to reduce tolerance to these metals in a tolerant strain of *S. meliloti* isolated from a mine (Lu et al., 2016, 2017). By contrast, in *Mesorhizobium metalladurins*, regulation of Zn in a few strains that were isolated from a heavy metal contaminated site appears to be due to enhanced *cis*-promoter driven regulation of the P_II_-type ATPase, cadA to achieve high tolerance (Maynaud et al., 2013, 2014). However, because of the correlated up-regulation of the adjacent genes in Mer operon in the Hg-tolerant Almadén strains, including hypothetical genes with no homology to known Hg transporters, there is the potential to assess incremental losses in tolerance by doing knock-outs of these genes. Such studies would identify the role of each gene linked within the operon, and the value of their up-regulation in connection with a tolerance phenotype. To summarize, the complete Mer operon is necessary for hypertolerance based on the strains we have isolated from Almadén, but the syntenic and structural diversity (variation in gene order and presence/absence variation) among α-proteobacteria, combined with extremely high upregulation of the Mer operon genes suggest that *cis*-regulation within the operon also may also play a very large role in achieving the level of tolerance that we observe in the collection of rhizobia from Almadén (i.e., MIC up to 250 μM of Hg) which may have evolved further due to exposure and contact with Hg in this highly toxic environment. We therefore speculate the operon likely evolved by a succession of horizontal gene transfer followed by *cis*-regulatory evolution of Mer genes.

## Conclusion

Despite the presence of gene annotations in publicly available genome data that show the presence of Mer operons in nitrogen-fixing bacteria such as *Mesorhizobium*, *Rhizobium* and *Sinorhizobium*, we presented the first experimental evidence of a functional Mer operon in nitrogen-fixing bacteria. Because the first line of defense against accumulation of toxic ions occurs in the plant roots, symbiotic rhizobia that can evacuate a toxic heavy metal like Hg from roots could be very beneficial because legumes are a staple food source for livestock and humans. The soil microbiota can play an important role in plant tolerance to these metals during symbiotic interactions with host-plants. Moreover, comparison of gene expression between non-Hg tolerant and the Hg-tolerant strains in nodules suggests there is little cost to possessing a Mer operon and the Hg-tolerant strain showed upregulation of important genes that would positively affect nitrogen-fixation for the host-plant. Furthermore, plants grown on contaminated soils may accumulate heavy metals in aerial parts such as leaf tissues and seeds and can result in severe health consequences for foraging animals and humans if these metals enter the food supply (Peralta-Videa et al., 2009). The presence of a Hg tolerance adaptation in symbiotic α-proteobacteria strains which can be used to fix nitrogen for host plants, improve nitrogen in soils at the entire plant community/population scale, while also contributing to heavy metal detoxification, may be an advantage to future efforts in removing toxins from plants (Narayanan and Ma, 2023), and also in bioremediation using legume-rhizobia interactions (Fagorzi et al., 2018).

## Methods

### Isolation and growth conditions of rhizobia

Rhizobia used in this study were isolated from nodules of wild plants that were collected from the Almadén mining region in Spain (See Nonnoi et al., 2012; Ruiz-Díez et al., 2012; 2013 for specific information regarding each strain). Originally, 33 *Sinorhizobium medicae* and 26 *Rhizobium leguminosarum* bv. *trifolii* strains were isolated from nodules of *Medicago* and *Trifolium* species respectively, which were growing at different locations near the abandoned Almadén mercury mine in Spain. Their tolerance was assessed using minimum inhibitory concentrations (MIC) over ranges of 25-250 μM. Among these, we selected 9 *S. medicae* and 5 *R. leguminosarum* strains (Table S1) that showed a broad range in Hg tolerance. Rhizobia strains were cultivated in liquid medium at 28°C in a shaking incubator until they reached an OD_600_ of 0.7. Total DNA was extracted using the Machery-Nagel NucleoSpin Microbial DNA Mini kit for DNA from microorganisms (Item number: 740235.50). During the procedure, bacteria cells were lysed using a Qiagen TissueLyser bead mill for 5-8 minutes.

### Genome sequencing, assembly, and annotation

Genomic sequence data were generated using a Pacific Biosciences (PacBio) RS II sequencer at the University of Minnesota Genomics Center with one PacBio single molecule real time (SMRT) cell per strain. Genomes were assembled using hgap version 4.0 (Chin et al., 2013) in the SMRTlink Server software (version 7) with default assembly parameter, using a similar pipeline described in Nelson et al., (2018). Subsequently, the genomes were circularized and contigs were verified using Circulator version 1.5.5 (Hunt et al., 2015) https://github.com/sanger-pathogens/circlator. To annotate genomes we used RASTtk https://rast.nmpdr.org (Brettin et al., 2015) using default settings. For the strains used for RNAseq, we also annotated the genome using the NCBI Prokaryotic Genome Annotation Pipeline (PGAP https://www.ncbi.nlm.nih.gov/genome/annotation_prok/) (Tatusova et al., 2016)in order to use these annotations to link GO-terms for GO-enrichment analysis. The annotations used throughout this manuscript are based on RASTtk, as this method was able to identify mercuric ion reductases better than PGAP. Protein sequences that were used in subsequent phylogenetic analyses and running OrthoFinder (Emms and Kelly, 2019) to label homologous genes used the RASTtk annotations, but we included PGAP annotations of strain *S. medicae* AMp08 in the Orthofinder analysis to link all annotations to a common strain. The AMp08 strain was used because it has a Mer operon on an accessory plasmid and the genome was completely assembled and circularized into four complete chromosomes and plasmids.

### Synteny analysis of whole genomes and Mer operon and phylogenetic analysis of merA homologs

To check for synteny between focal strains with previously published reference strains of *S. medicae*, we used Sibelia 2.2.1 (Minkin et al., 2013) using the command ./Sibelia -s loose -q. We aligned the two Hg tolerant Almadén strains AMp08 and SMp01 strains to *S. medicae* WSM419 to assign the chromosomes and pSym plasmids according to (Reeve et al., 2010a). For the *R. leguminosarum* strain Stf07, we aligned the *R. leguminosarum* WSM1325 genome to assign chromosomes and accessory plasmids (Reeve et al., 2010b). We used OrthoFinder (Emms and Kelly, 2019) to cluster genes in order to obtain orthogroups that can connect the genes from each independent annotation in this study and other publicly available genomes. For the synteny analyses, we created orthologue tables using OrthoFinder (Emms and Kelly, 2019). IMG annotations (cog, ko and pfam) were mapped onto the orthologue tables, and each gene annotation contains the gene start, end and strand direction information for each gene. Additionally, we used ortho groups to assign a consensus annotation when a given protein was not annotated as 100% the same annotation or to fill genes of unknown function when a consensus annotation was available. Note, these situations are rare, but the orthologue group clustering helps to make sense of either missing or misannotated genes. To produce the synteny plot displayed in Figure S4, we used the R package gggenomes (https://github.com/thackl/gggenomes). We could find MerA homologs that possessed flanking MerT, MerC, MerP and MerB genes for the synteny analysis of the Mer operons in our dataset. We also built two separate phylogenies within this pipeline, individual gene trees, and species trees based on a set of 56 concatenated markers from 72 bacterial genomes. To produce the gene trees, we aligned the markers using mafft (Katoh and Standley, 2013) then produced a tree with FastTree2 (Price et al., 2010), that was used to orient the Almadén genomes with others in IMG to make a circular phylogeny that represented relationships of α-proteobacteria (Figure S1). Mercuric acid reductase A (merA) homologs were identified using Orthofinder, and we aligned amino acid sequences of the four homologs merA1 (OG0004493), merA2 (OG0001010) merA2 and dihydrolipoamide 2-oxoglutarate dehydrogenase (D2OD) (OG0000957), and MerA (OG0009124) using Geneious and Clustal. We used Blast to obtain outgroup sequences, many of which were annotated as MerBA in NCBI/GenBank. We used Mr. Bayes v3.2.7 on the CIPRES webserver (https://www.phylo.org/index.php) and sampled from 500,000 trees to obtain a consensus tree with posterior probability scores.

### Modeling the protein structure of mercury reductase A homologs

Protein structures were modeled through Google ColabFold with MSA mode MMseqs2, pair mode unpaired+paired, and other default settings (https://colab.research.google.com/github/sokrypton/ColabFold/blob/main/AlphaFold2.ipynb). ColabFold was developed from AlphaFold2 for accelerated structure prediction of proteins and their complexes, and it matches AlphaFold2 and AlphaFold Multimer in prediction quality of tested datasets (Mirdita et al., 2022). Modelled structures were superposed with COOT (Emsley and Cowtan, 2004) for the structural comparison. PyMOL (Schrödinger, LLC, 2015) was used to visualize the structures and prepare figures.

### Isolation of RNA for rhizobia grown under free-living conditions

We selected four *S. medicae* strains which show the highest and lowest tolerance to Hg in our collection and also had high quality reference genome assemblies for transcriptomics analysis. In all experiments in this work, HgCl_2_ has been used as the source of Hg. From here on, Hg tolerance, sensitivity and stress, refer to HgCl_2_. *S. medicae* strains AMp08 and SMp01 (Hg-tolerant), AMp07 and VMo01 (Hg-sensitive) were grown in liquid TY medium in the absence or presence of 4 μM HgCl_2_ at 28°C in a shaking incubator until they reached an OD_600_ of 0.8. Subsequently, we directly added the Qiagen RNAprotect Bacteria Reagent to the bacterial culture in a 1:2 ratio (culture: RNA stabilizer). The mixtures were vortexed for 5 s and incubated for 5 min at room temperature (23°C), then centrifuged at 5,000 × g for 10 min at 20°C. Pellets were frozen in liquid nitrogen and bacterial lysis was performed by resuspension and incubation of the cell pellet in 1 mg/ml lysozyme from chicken egg whites (Sigma-Aldrich) in Tris-EDTA buffer, pH 8.0. Total RNA was extracted using the Qiagen RNeasy Mini Kit using the manufacturer’s conditions specified.

### Isolation of RNA from nodules

To evaluate the effects of Hg on *S. medicae* gene expression inside of nodules of *M. truncatula* host-plants, we grew two *M. truncatula* genotypes (HM302, HM304) in sterilized turface as a soil substrate, then inoculated the plants with two *Sinorhizobium medicae* strains (AMp07, AMp08) that have different Hg-tolerance. Plants were well-replicated (average of 20 plants per assay) and completely randomized. Prior to planting, seeds were treated in concentrated sulfuric acid for 3 minutes and washed 5 times with MiliQ water. Then seeds were surface sterilized using 50% bleach with 0.1% Tween20 for 3 minutes and washed 8-10 times with sterilized MiliQ and left in sterile water for 2 hours at room temperature in dark. The seeds were placed on sterilized filter paper on petri dishes and kept overnight at 4°C in dark. Later the petri dishes were kept in growth chamber at 21°C, 40% humidity, on a 16-h-light/8-h-dark photoperiod for a week, then seedlings were transferred to the sterilized Turface:vermiculite (2:1) soil (LESCO-turface all sport pro soil) in plant growth chamber. Plants were cultivated in a walk-in growth chamber in 16-h-light/8-h-dark photoperiod and 22°C conditions and watered every 5-7 days with sterilized ½ strength of B&D medium with low concentration of KNO_3_ (0.5mM). Prior to inoculation, KNO_3_ was not added while watering the seedlings. Seedlings were inoculated with *S. medicae* AMp07 and AMp08 rhizobia strains 5-7 days after being transplanted to the sterilized turface soil. The inoculum was prepared by culturing rhizobia at 28°C in sterilized liquid TY medium until an OD600 of 0.8 was reached. Subsequently, the culture was centrifuged at 4000 rpm for 5 mins and the supernatant removed and washed 2 times with autoclaved 0.9% NaCl. The pellet was resuspended in 0.9% NaCl for inoculation. 21 days post inoculation (21dpi), the plants were treated with 100μM HgCl_2_, while non-treated plants were watered with ½ strength of B&D medium supplemented with 0.5 mM concentration of KNO_3_. After 7 days of treatment, the nodules were harvested and quickly frozen in liquid nitrogen and stored at −80 °C for RNA-seq.

### Library preparation and RNAseq of rhizobia grown under free-living conditions and in nodules

Library preparation and sequencing of the free-living rhizobia samples was done at the DOE Joint Genome Institute under Community Science Project (Functional Genomics) 506402. RNAseq data for each library was generated using the Illumina NovaSeq S4 platform which generated 2x151 bp reads. BBDuk (version 38.90) was used to remove contaminants (BBDuk, BBMap and BBMerge commands used for filtering are placed in file: 52510.2.364064.AAGAAGGC-AAGAAGGC.filter_cmd-MICROTRANS.sh), trim reads that contained adapter sequence and homopolymers of G’s of size 5 or more at the ends of the reads, right quality trim reads where quality drops below 6, remove reads containing 1 or more ‘N’ bases, remove reads with average quality score across the read less than 10, having minimum length <= 49bp or 33% of the full read length.

For nodule samples, construction of the RNAseq libraries and sequencing on the Illumina NovaSeq 6000 were performed at the Roy J. Carver Biotechnology Center at the University of Illinois at Urbana-Champaign. Purified DNAsed total RNAs were run on a Fragment Analyzer (Agilent) to evaluate RNA integrity. The total RNAs were converted into individually barcoded RNAseq libraries with the Universal Plus mRNA-Seq Library Preparation kit from Tecan, using custom probes against *Medicago* and *S. melliloti* rRNA designed by Tecan. Libraries were barcoded with Unique Dual Indexes (UDI’s) which have been developed to prevent index switching. The adaptor-ligated double-stranded cDNAs were amplified by PCR for 10 cycles. The final libraries were quantitated with Qubit (ThermoFisher) and the average cDNA fragment sizes were determined on a Fragment Analyzer. The libraries were diluted to 10nM and further quantitated by qPCR on a CFX Connect Real-Time qPCR system (Biorad) for accurate pooling of barcoded libraries and maximization of number of clusters in the flowcell. The barcoded RNAseq libraries were loaded on one SP lane on an Illumina NovaSeq 6000 for cluster formation and sequencing. The libraries were sequenced from both ends of the fragments for 150bp from each end. The fastq read files were generated and demultiplexed with the bcl2fastq v2.20 Conversion Software (Illumina).

### Mapping of RNAseq reads and differential expression analysis

Free-living rhizobia RNAseq reads from each *S. medicae* strain were mapped to the newly assembled genomes specific to each strain. For nodule samples, concatenated reference genomes of *Medicago truncutala* v5.0 and either *S. medicae* AMp07 or AMp08 were used for mapping the dual-transcriptomics samples. Indexing of each genome was done using STAR v2.7.10a (Dobin et al., 2013). FastQC v0.11.9 (Andrews, S, 2010) and Trimmomatic v0.39 (Bolger et al., 2014) were used for quality control and adapter removal with default parameters. The resulting high-quality reads were mapped to the appropriate reference genome assembly using STAR v2.7.10a and raw counts were generated using the GeneCounts feature. Differentially expressed genes were identified using DESeq2 v1.34.0 and p-values were adjusted using Benjamini and Hochberg’s correction (Love et al., 2014) PhyloFlash v3.4 (Gruber-Vodicka et al., 2020) was used to check for contamination of samples using bacteria databases to align raw data. To aggregate read counts for each gene, we used a custom python script aggregate_counts.py (https://github.com/mclear73/RNAseq_Miniworkshop/blob/main/Processing%20Raw%20RNAseq%20Data%20into%20Counts.md) which generates a count table and metadata file to be used for input for DESeq2, which we used to calculate differential expression in Hg-treated samples compared with control samples. Genes with an adjusted p-value < 0.05 and log_2_ fold-change ≥ 1 were considered differentially expressed. For direct comparison of gene counts between samples, raw gene counts were normalized using the trimmed mean of m-value (TMM) method (Robinson and Oshlack, 2010) from edgeR v3.36.0 (Robinson et al., 2010). Transcripts per kilobase million (TPM) were calculated using a custom python function.

### Ortholog Comparison and GO-Enrichment Analysis

Orthogroups were identified for 20 related bacterial strains including AMp08 annotated with the Prokaryotic Genome Annotation Pipeline (PGAP) (Tatusova et al., 2016) with OrthoFinder v2.5.4 (Emms and Kelly, 2019) with default parameters. The complete genome annotation for AMp08 was generated using NCBI PGAP (v2022-02-10.build5872) because the RAST annotations cannot be directly linked to GO-terms. To allow for a standardized comparison of enriched GO-terms between strains, all GO enrichments were performed with the PGAP annotation of AMp08 via orthologs determined by OrthoFinder. Using custom python scripts, DEGs were mapped to the Orthogroup provided by Orthofinder and all AMp08 genes within the Orthogroup were used as input for GO Enrichment. Enriched GO-terms for DEGs were identified using a custom python script for g:Profiler2 using the built in “g_SCS” function that corrects for multiple tests (Raudvere et al., 2019) Tests for overrepresentation were performed for the biological processes (BP), Cellular components (CC), and molecular function (MF) databases based on the PGAP annotation for AMp08.

### Transfer of plasmid containing Mer operon into non-tolerant strains

The *Sinorhizobium medicae* strain AMp08 from the Almadén mine is highly tolerant to Hg harbors a Mer operon on plasmid. To isolate the plasmid, the AMp08 strain was grown in liquid TY medium (28°C, 220 rpm) until OD_600_ = 0.8 was reached. Afterward the plasmid was isolated using GenElute plasmid miniprep kit (Sigma Aldrich). The recipient strains (AMp07 and WSM419) were made electrocompetent using the protocol described in (Garg et al., 1999). Specifically, the strains were grown in liquid TY medium at 28°C for 2 days at 220 rpm until they reached mid-logarithmic stage (OD_600_ = 0.8-1.0). Subsequently the cells were chilled on ice for 15 to 30 minutes, and then collected from the pellet after centrifugation at 3700 g for 10 minutes at 4°C. The cell pellet was washed four times with chilled deionized water, then with 10% glycerol. The cells were resuspended in 10% glycerol and stored at -80°C in 100 μl aliquots. For electroporation, 4 μl (∼250-300 ng) of AMp08 plasmid DNA was mixed with the recipient strain by gentle vortexing and kept on ice for 30 minutes. The cell-DNA mixture was then loaded into a pre-chilled electroporation cuvette with a 0.1-cm gap and was exposed to a single pulse of high voltage of field strength 14 kV/cm for 5.8 ms. After the electroporation, the cuvettes were kept on ice for an additional 10 minutes. Once the cells were allowed to grow in liquid TY media for an additional 1 hour, the cell culture was plated on TY plates containing 200 and 250 μM HgCl_2_ to select for transformed strains. Finally, to confirm the positive transformation, genomic DNA was isolated from the positive colonies and the presence of AMp08 MerA gene was confirmed using PCR. To confirm if the Mer operon was functional in the transformed strains, qPCR was conducted to check the expression levels of key Mer operon genes (MerA and MerT) as well as merA1, which is expressed in all the *S. medicae* strains. Briefly, RNA was isolated using the methodology described above. 1 μg of total RNA was used to synthesize cDNA using SuperScript III cDNA synthesis kit (Invitrogen, U.S.A) with random hexamer primers, and the cDNA samples were diluted 1:10 in dH2O. qRT-PCR assays were conducted using gene specific primers (Supplementary table X. Primer list) using SYBR green master mix (Bio-Rad). The reaction was run for a total of 40 cycles, and the relative expression was calculated using the comparative Ct method (2 ^-ýýCt^) method. The threshold cycle (Ct) value of the reference gene, namely 16S rRNA was used to normalize the transcript levels of all genes. All the results shown here include data averaged from three biological replicates, which in turn included three technical replicates each.

### Quantifying minimum inhibitory concentration (MIC)

All the rhizobia strains used in this study (AMp07, AMp08, WSM419) were grown in autoclaved liquid TY medium until an OD_600_ of 0.8 was reached. To quantify MIC, a stock solution of TY medium and 1% agar was prepared by adding 100 mM of filter sterilized HgCl_2_ stock solution to autoclaved TY to get the required concentrations containing 0 μM, 25 μM, 50 μM, 75 μM, 100 μM, 150 μM, 200 μM, 250 μM HgCl_2_. The cultures were serially diluted 1:5, 1:10, 1:100 using TY solution and 10 μl each of the non-diluted and diluted cultures were added to all the Hg-containing TY plates. Subsequently, the plates were incubated at 28°C for 2 days and images were taken. The MIC represents the lowest concentration of Hg at which the growth of a strain is inhibited, and it was confirmed for all the original strains (AMp07, AMp08, WSM419) and determined for the engineered AMp07 and WSM419 strains.

## Data Accessibility

Raw data for genome assemblies (PacBio reads) https://www.ncbi.nlm.nih.gov/bioproject/?term=PRJNA565470

Raw data for free-living rhizobia RNA-seq (150 bp paired end Illumina reads) https://www.ncbi.nlm.nih.gov/bioproject/?term=PRJNA996196

Raw data for rhizobia reads from dual-transcriptomics (150 bp paired-end Illumina reads) https://www.ncbi.nlm.nih.gov/bioproject/?term=PRJNA986885

## Acknowledgments

We would like to thank Katy Heath, Alvaro Hernandez and Chris Wight at the University of Illinois at Urbana-Champaign for consulting, oligo design, and rRNA cleanup and sequencing for the nodule samples. We thank Sanhita Chakraborty at the Institute for Advancing Health through Agriculture at Texas A&M for useful comments and interpretation of data. Funding for this project came from the Department of Energy Quantitative Plant Science Initiative SFA, USDA-ARS Project Number 3092-53000-001-000D, the Joint Genome Institute for RNAseq of the free-living rhizobia under Community Science Project (Functional Genomics) 506402 (Award DOI: 10.46936/10.25585/60001331) awarded to TP, and the Agencia Estatal de Investigación, Spain, grant number PID2021-125371OB-I00 awarded to JJP. The work conducted by the U.S. Department of Energy Joint Genome Institute (https://ror.org/04xm1d337), a DOE Office of Science User Facility, is supported by the Office of Science of the U.S. Department of Energy operated under Contract No. DE-AC02-05CH11231.

