## SupplementaryFigures for "Local adaptation to mercury contamination by nitrogen-fixing rhizobia is driven by horizontal gene transfer, copy number, and enhanced gene expression"

**Supplementary Figures.**

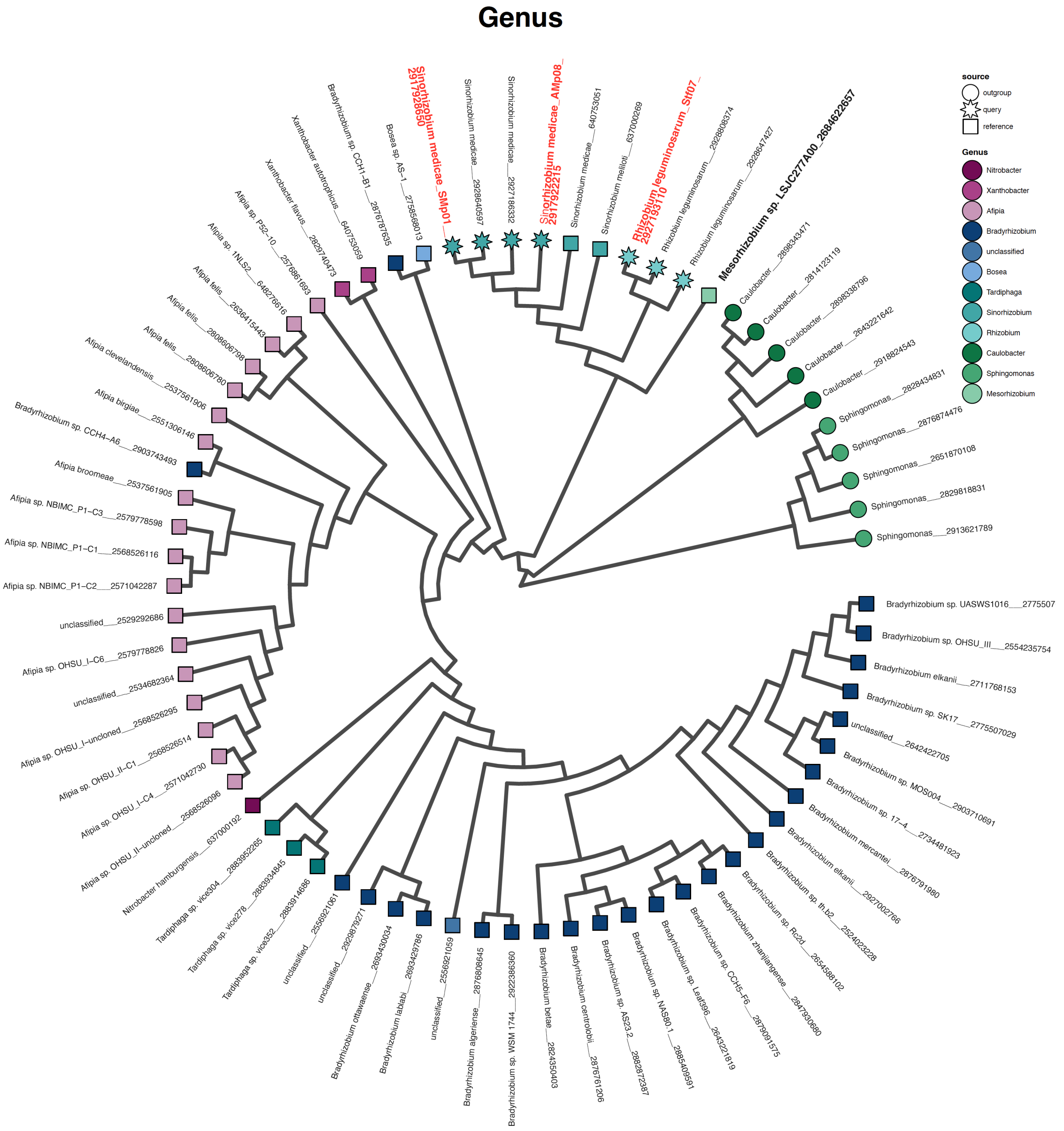


Figure S1. Phylogenomic analysis of α-proteobacteria genera and species that were used to identify syntenic genomic regions and to verify the expected phylogenetic relationships of focal strains from Almadén. The three Almadén strains, *Sinorhizobium medicae* AMp08, SMp01 and *Rhizobium leguminosarum* Stf07 which are the most Hg-tolerant strains are labeled in red. *Mesorhizobium* sp. LSJC277A00 (bold) has a Mer operon.

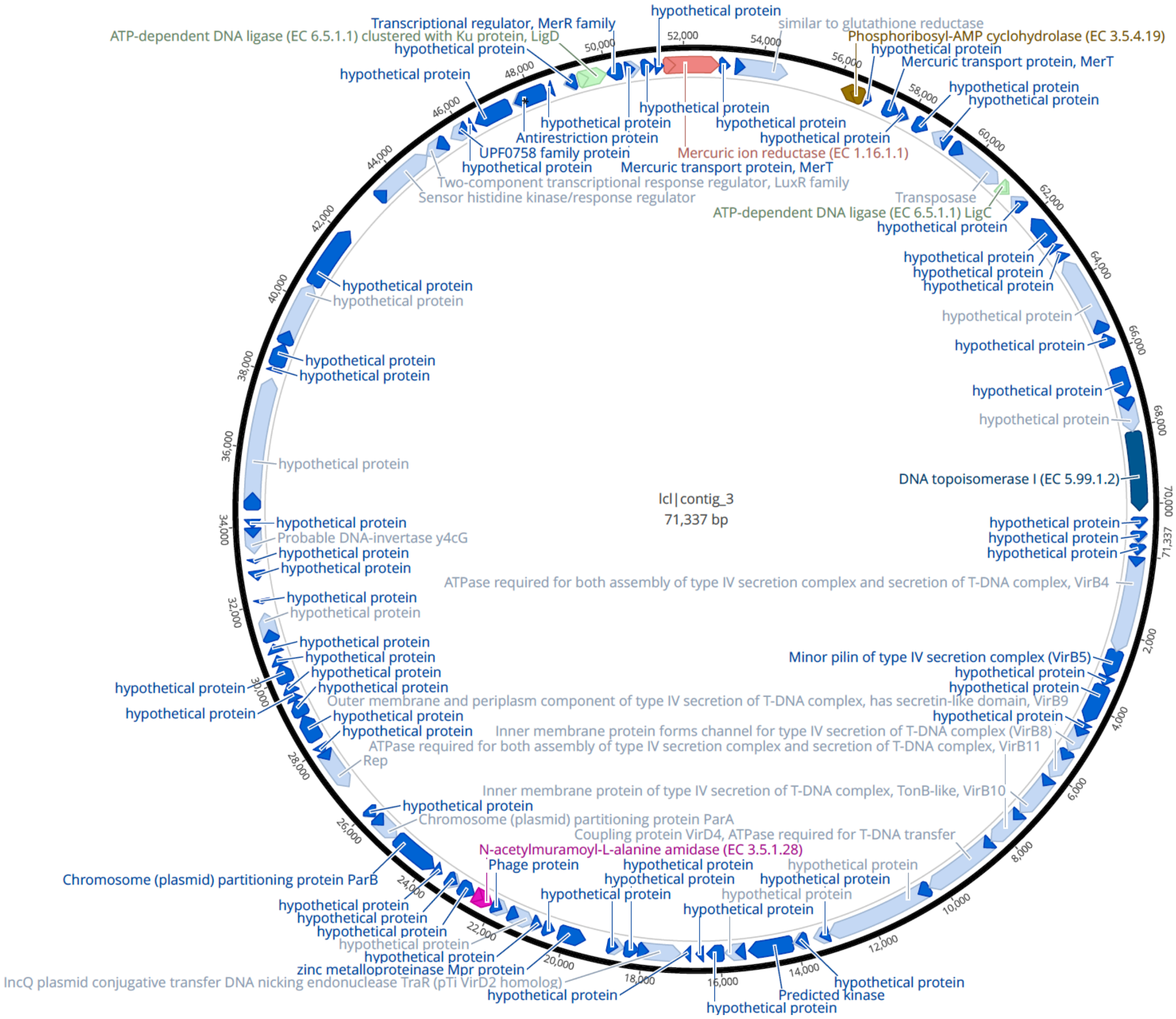


Figure S2. The accessory plasmid containing the Mer operon in strain AMp08 is 71 kb and contains many uncharacterized genes (hypothetical proteins) and shows no homology or synteny with WSM419 (see Figure 1a).

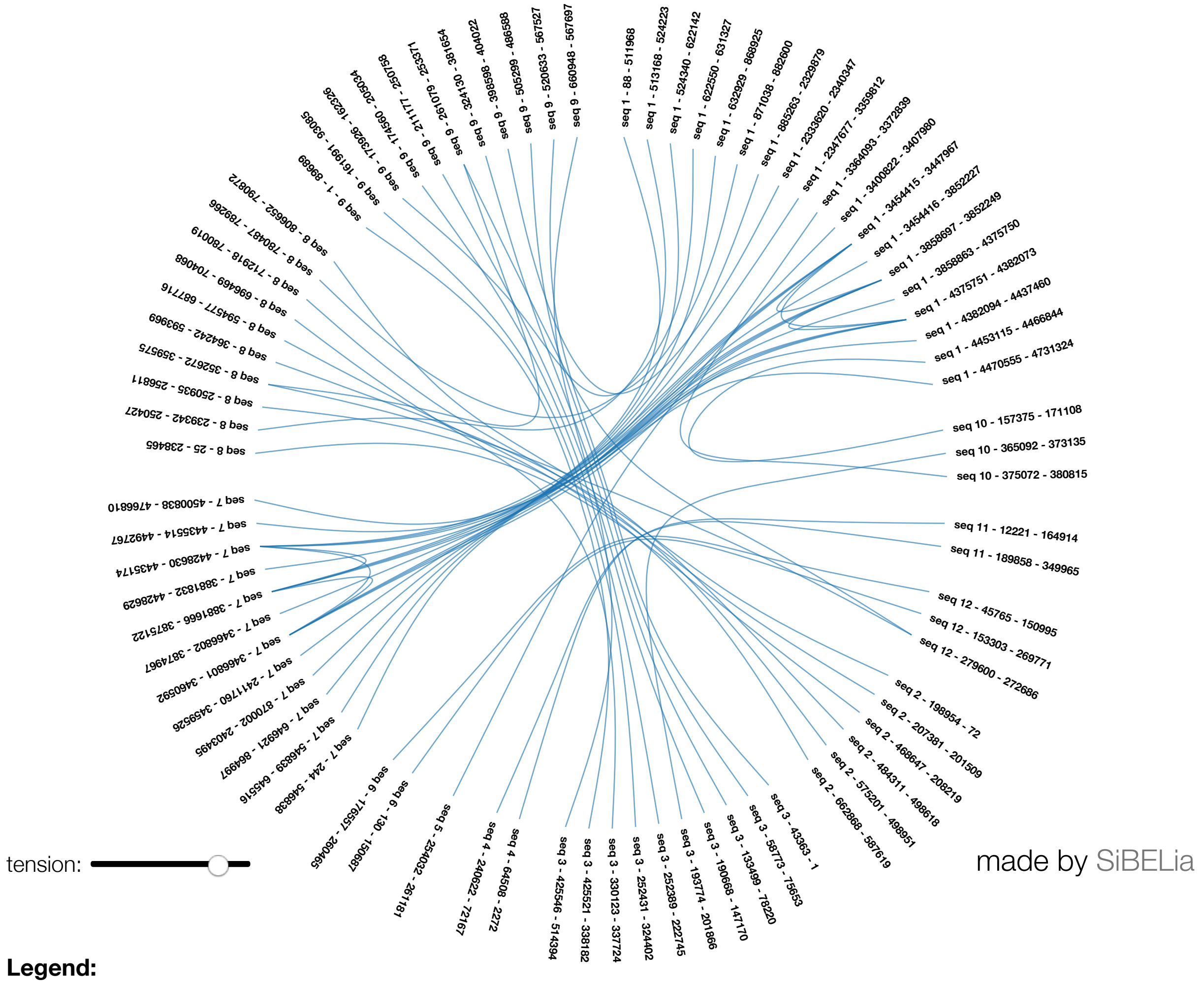


**Legend:**

seq 1 : Stf07_contig_0
seq 2 : Stf07_contig_1
seq 3 : Stf07_contig_2
seq 4 : Stf07_contig_3
seq 5 : Stf07_contig_4
seq 6 : Stf07_contig_5
seq 7 : NC_WSM1325_012850.1
seq 8 : NC_WSM1325_012848.1
seq 9 : NC_WSM1325_012858.1
seq 10 : NC_WSM1325_012853.1
seq 11 : NC_WSM1325_012852.1
seq 12 : NC_WSM1325_012854.1

Figure S3. Whole genome synteny of *Rhizobium leguminosarum* strain STf07 with *R. leguminosarum* WSM1325. The chromosome of WSM1325 is NC_012850.1 Others are plasmids: pR132501=NC_012848.1, pR132502=NC_012858.1, pR132503 = NC_012853.1 pR132504 =NC_012852.1 pR132505 = NC_012854.1. Genome alignment and circos figure was done using Sibelia v2.1.1.

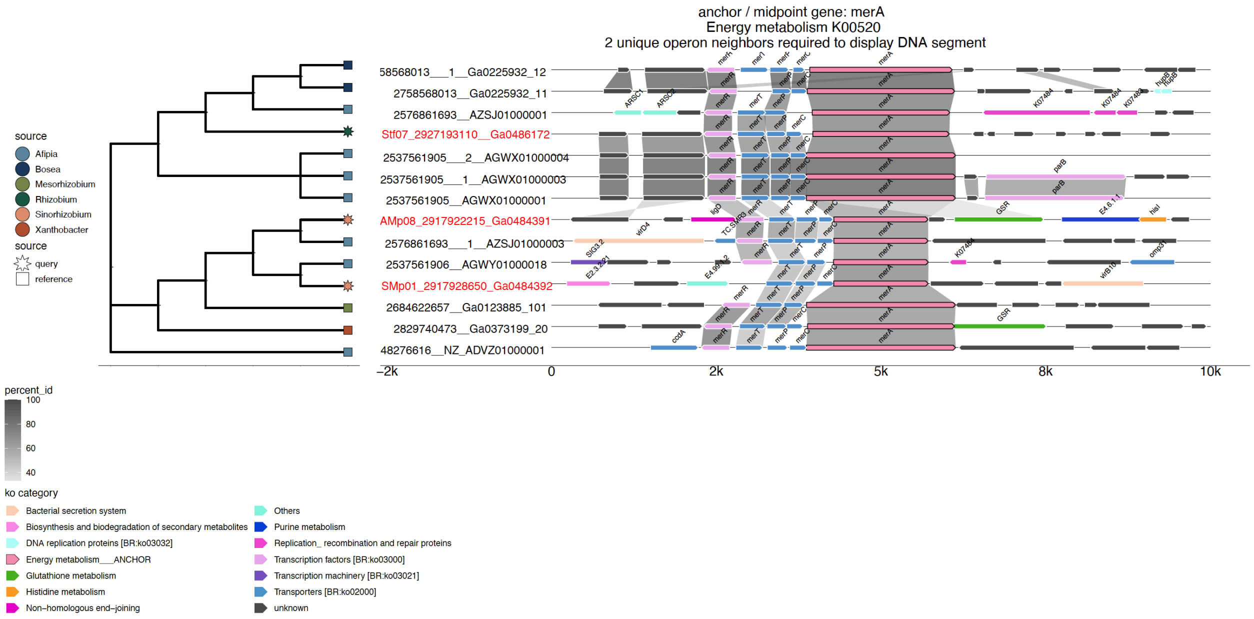


Figure S4. Synteny of three Almaden mine strains anchored at the merA gene as a query, with other α-proteobacteria in the IMG database (https://img.jgi.doe.gov). The phylogeny shows the three Almadén strains names with high tolerance to Hg are colored using red text, and were used as query sequences indicated by the star character at the tips of the phylogeny.

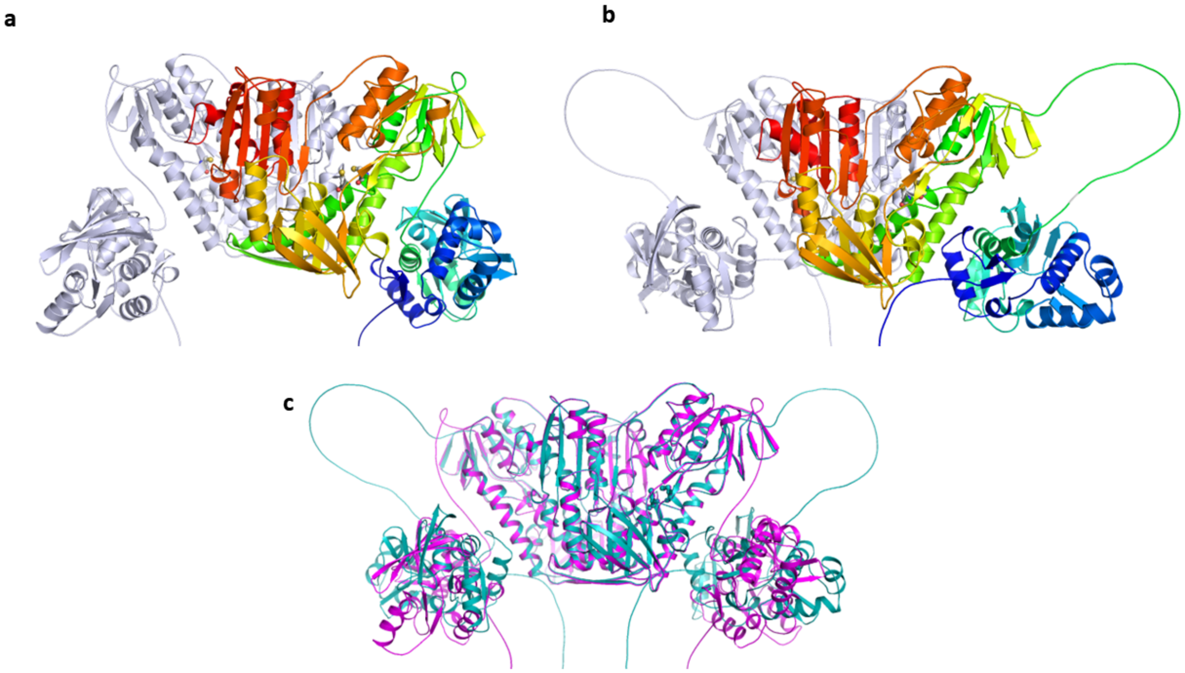


Figure S5. AlphaFold structures of dimeric MerBA fusion proteins from *R. leguminosarum* strain STf07 (a) and *Mesorhizobium* sp. LSJC277A00 (b). The two subunits of MerBA dimer are shown in grey and spectrum (Blue = N-terminus, Red = C-terminus) colors. (c). Overlay of the structures of *R. leguminosarum* strain STf07 (magenta) and *Mesorhizobium* sp. LSJC277A00 (cyan). Cysteine residues are shown in the stick representation.

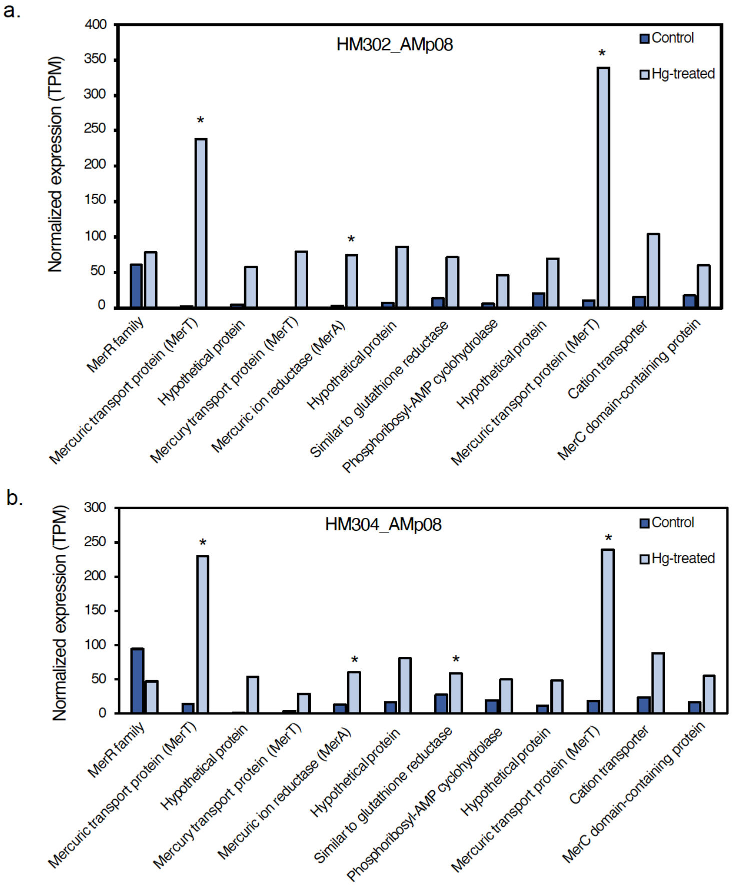


Figure S6. Expression of Mer operon genes in nodules in two different *M. truncatula* host plant genotypes, formed by the *S. medicae* strain AMp08 in control conditions and following Hg treatments. (a) The Mer operon gene expression in nodules of the HM302 host plant, which is a low Hg accumulating plant genotype. (b) The Mer operon gene expression in nodules of the HM304 host plant, which is a high Hg accumulating plant genotype. The host-plants inoculated with rhizobia were treated with either 0 µM $HgCl_2$ (Control) or 100 µM $HgCl_2$ (Hg-treated). Normalized expression levels are the mean TPM values of three biological replicates.

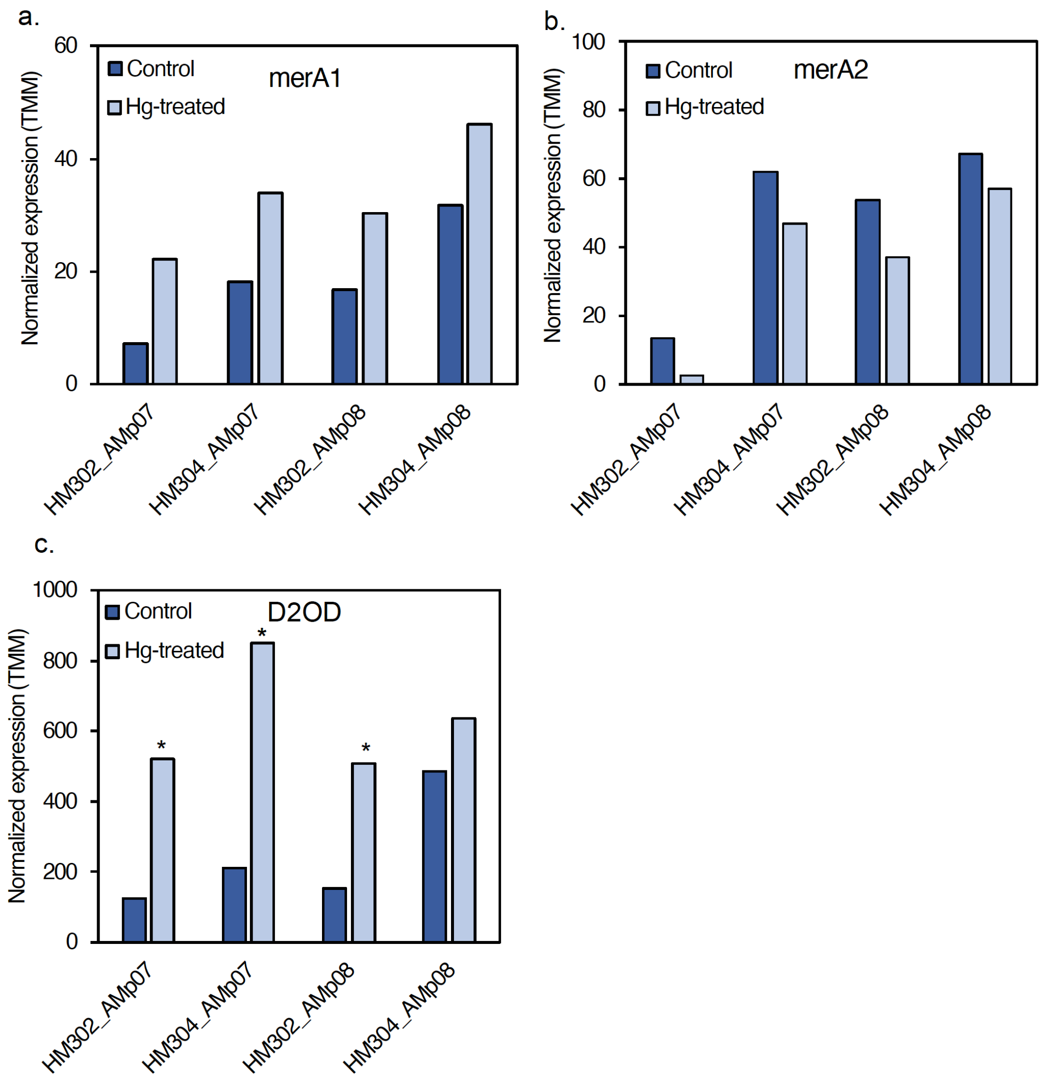


Figure S7. Expression of mercuric ion reductase gene homologs in rhizobia present in nodules in control and following Hg treatments. (a) Mercury ion reductase (merA1) shows no significant change in response to Hg treatment in any of strains (AMp08 and AMp07). (b) Mercury ion reductase (merA2) shows no significant change in response to Hg stress in any of the strains tested (AMp08 and AMp07). (c) Dihydrolipoamide 2-oxoglutarate dehydrogenase (D2OD) showed significant upregulation in all the strains tested (AMp08 and AMp07). The normalized expression levels are shown as the mean TMM values of three biological replicates.

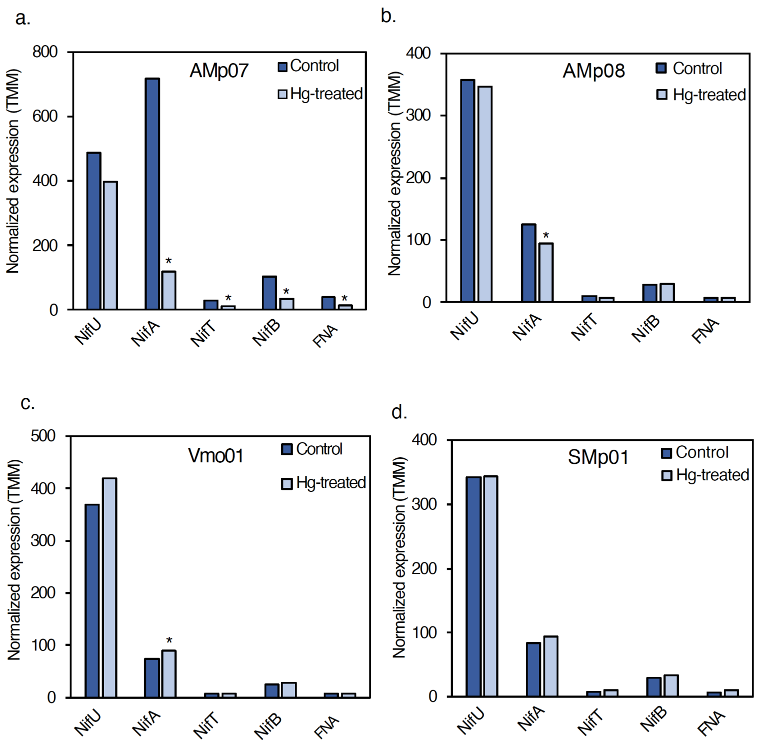


Figure S8. Expression of *nif* genes in free-living rhizobia in control and Hg treatment conditions. (a) In the non-tolerant strain AMp07, the expression levels of *nifU*, *nifA*, *nifT* and 4Fe-4S ferredoxin nitrogenase-associated gene (FNA) were significantly downregulated. A small but significant change in *nifA* expression was observed in AMp08 (b), and small but significant change in *nifA* expression was observed in VMo01 (c), and no significant changes in SMp01 (d). Normalized expression levels are reported as mean TMM values of three biological replicates.

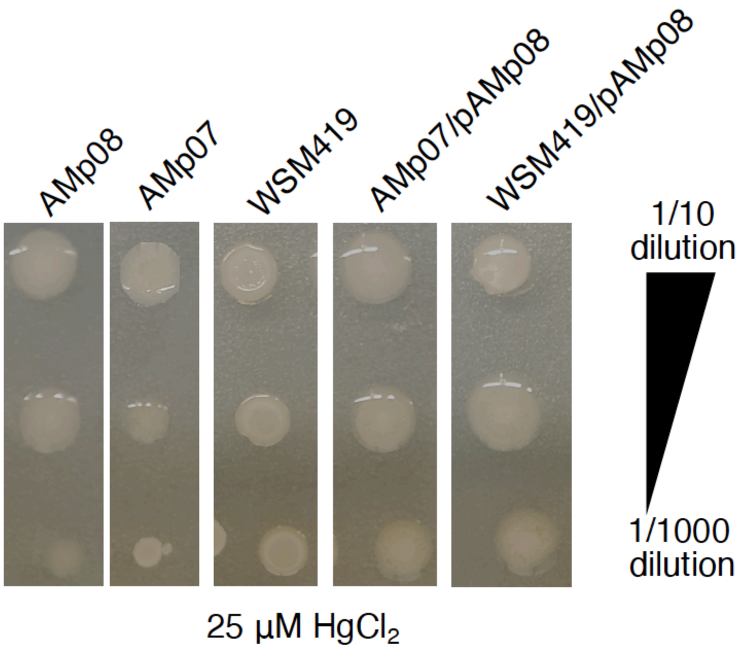


Figure S9. Minimum inhibition concentration (MIC) assay shows the basic Hg-tolerance in non-tolerant strains, the Hg- tolerant strain AMp08 and the two strains transformed with the accessory plasmid from AMp08 (AMp07/pAMp08; WSM419/pAMp08). Each MIC image represents strains (from left to right: AMp08, AMp07, WSM419, AMp07/pAMp08, WSM419/pAMp08) grown on plates containing 25 μM $HgCl_2$. Each strain was grown at three dilution factors (from top to bottom): 1/10, 1/100 and 1/1000.
